# Escape from X inactivation is directly modulated by levels of Xist non-coding RNA

**DOI:** 10.1101/2024.02.22.581559

**Authors:** Antonia Hauth, Jasper Panten, Emma Kneuss, Christel Picard, Nicolas Servant, Isabell Rall, Yuvia A. Pérez-Rico, Lena Clerquin, Nila Servaas, Laura Villacorta, Ferris Jung, Christy Luong, Howard Y. Chang, Judith B. Zaugg, Oliver Stegle, Duncan T. Odom, Agnese Loda, Edith Heard

## Abstract

In placental females, one copy of the two X chromosomes is largely silenced during a narrow developmental time window, in a process mediated by the non-coding RNA Xist ^1^. Here, we demonstrate that Xist can initiate X-chromosome inactivation (XCI) well beyond early embryogenesis. By modifying its endogenous level, we show that Xist has the capacity to actively silence genes that escape XCI both in neuronal progenitor cells (NPCs) and *in vivo*, in mouse embryos. We also show that Xist plays a direct role in eliminating TAD-like structures associated with clusters of escapee genes on the inactive X chromosome, and that this is dependent on Xist’s XCI initiation partner, SPEN ^2^. We further demonstrate that Xist’s function in suppressing gene expression of escapees and topological domain formation is reversible for up to seven days post-induction, but that sustained Xist up-regulation leads to progressively irreversible silencing and CpG island DNA methylation of facultative escapees. Thus, the distinctive transcriptional and regulatory topologies of the silenced X chromosome is actively, directly - and reversibly - controlled by Xist RNA throughout life.

## INTRODUCTION

Sex-specific gene expression results in a multitude of sexually dimorphic phenotypes including diseases ^3–6^. One source of such differential expression between XX and XY individuals is the increased expression of some X-linked genes in females, due to their escape from XCI^4,7,8^. Higher levels of X-linked gene products in females may contribute to sex-specific regulatory programs and account for sex-biased disease predisposition, as shown in the case of autoimmune diseases and immune responses ^9–11^.

While the mechanism underlying the onset of X-chromosome wide gene silencing driven by Xist RNA is increasingly clear, the mechanisms that underlie escape from XCI and the eventual role of Xist have remained obscure ^12^. During early development, Xist RNA spreads *in cis* and coats one of the two X chromosomes leading to its almost complete transcriptional repression via factors such as SPEN ^2,13–17^. How a subset of “escapee” genes avoid or override this silencing, to be biallelically expressed from both active (Xa) and inactive (Xi) X chromosomes is not known ^18,19^. A small percent of X-linked escapees (∼3-7% in mice and ∼4-11% in humans^19–22)^) can evade silencing from the very onset of XCI. Such constitutive escape is found in most cell types and individuals and is conserved across different species ^12,19^. By contrast, a larger subset of X-linked genes (at least 20% in human and mice) show variable “facultative” escape, with genes becoming re-expressed following silencing in some tissues, and often variably between individuals ^23–28^. There is increasing evidence that the expression levels of escapees may have a profound impact on sexual dimorphism during development and in sex-biased disease predisposition ^4,29,30^.

Here we set out to test whether Xist plays any role in regulating escapees. Xist RNA is essential to trigger XCI during early development ^31–33^ and the prevailing view has been that it becomes dispensable later on for XCI maintenance in differentiated cells, where multiple layers of epigenetic factors such as DNA methylation, can lock in the silent state of the Xi ^34–37^. In the mouse, Xist is indispensable for the initiation of two waves of XCI ^12^, the first being imprinted (iXCI), leading to silencing of the paternally-inherited X chromosome (Xp) by E3.5 ^23,28,38,39^. The second wave of XCI is random (rXCI) and occurs in the embryo proper at around implantation (∼E5.5) ^28^. Random XCI can be faithfully recapitulated *in vitro* upon differentiation of mouse embryonic stem cells (mESCs) or by ectopic Xist induction in undifferentiated cells ^25,40–42^. Work on Xist tetracycline-responsive transgenes in mouse ESCs defined a critical time window during early differentiation that marked a shift from Xist-dependent gene silencing to Xist-independent and irreversible XCI, as well as the loss of Xist RNA’s capacity to initiate chromosome-wide gene silencing *de novo* ^33^. Recent work has shown that Xist RNA initiates XCI in early embryos or ESCs by recruiting its interactor SPEN to the Xi via the A-repeat domain at its 5’ end ^2,33^. Xist RNA also recruits (via hnRNPK) the Polycomb Repressive Complex 1 (PRC1), which is responsible for the Xi-accumulation of H2AK119ub which in turn recruits Polycomb Repressive Complex 2 (PRC2) ^2,43–45^. These repressive epigenetic changes are thought to mediate early maintenance of the inactive state and the accumulation of DNA methylation at CpG islands on the Xi is thought to be the key epigenetic change that fully locks in the maintenance of XCI in differentiated somatic cells ^46,47^.

However, there is increasing evidence, thanks to the use of highly sensitive approaches such as RNA FISH and allelic mRNA detection, that reactivation of silenced Xi genes, or upregulation of gene already partially escaping from XCI, can in fact occur upon *Xist* loss and that this may impact tissue homeostasis and even contribute to disease ^37,48–53^. In these studies, genes that show reactivation upon Xist loss often correspond to genes that have been shown to escape in other contexts, tissues or cell types ^37,48–52,54^. Both in humans and mice, deletion or depletion of *XIST*/*Xist* leads to de-dampening of escapee expression levels from the Xi, where these genes are generally transcribed at lower levels compared to the Xa ^37,48,49,52^. Thus, *Xist* may actually participate in X-linked gene silencing in adult cells, contrary to previous thinking that its main role is only the initiation of XCI. Nevertheless, the exact causality between *Xist* RNA levels, the transcriptional activity of escapees, and the factors involved in Xist’s silencing function in differentiated cells is not clear. These are important questions, as over- or under-expression of X-linked escapees can impact development and disease.

## RESULTS

### Increased Xist RNA levels can silence Xi escapee genes in differentiated cells

To explore the role that Xist RNA levels might play in directly modulating the expression levels of escapees in post-XCI differentiated cells, we assessed whether expression levels of escapees correlate with Xist RNA levels in 21 female NPCs F1 hybrid lines (*Mus musculus domesticus* (129/Sv) x *Mus musculus castaneus* (Cast/Eij))^55^. We first characterised the XCI status in these 21 clones by performing transcriptomic analysis [Extended Data Fig. 1a,b,c]. For each informative X-linked gene we used allele-specific RNA quantifications to calculate its allelic ratio (reads^Xi^/[reads^Xi^+reads^Xa^]) ^56^. X-linked genes displaying biallelic expression (allelic ratio of greater than 0.1), were considered escapees (i.e. Xi-specific expression levels contribute to at least 10% of the gene’s total expression) [Extended Data Fig. 1c]. We confirmed substantial variability across different NPC clones in the numbers of escapees per clone, ranging from 48 to 124 genes escaping XCI out of 379 X-linked genes for which we have allelic information, as previously shown [Extended Data Fig. 1d] ^56,57^ and also consistent with human inter-individual data ^4,7,20–22,58^. We assessed whether higher levels of Xist RNA correlate with lower expression levels of escapees chromosome-wide and found a significant negative correlation (p=0.012, correlation test), suggesting that Xist may indeed play a role in tuning the dosage of X-linked escapees on the Xi [Fig. 1a].

To test this directly, we established a system in which we could systematically test the impact of increased Xist RNA levels on the transcriptional activity of escapees in post-XCI NPC cells. We generated clonal NPC lines by *in vitro* differentiation of female TX1072 mESCs ^59^, which is a polymorphic F1 hybrid line (*Mus musculus castaneus* (Cast/Eij) x *Mus musculus domesticus* (C57BL/6)) carrying a doxycycline (Dox) inducible promoter upstream of the *Xist* endogenous locus on the C57BL/6 X chromosome ^59,60^. TX1072 ESCs undergo random XCI during differentiation in the absence of Dox. We thus generated (without Dox) a heterogenous pool of NPCs in which either the C57BL/6 (B6) or the Cast/Eij (Cast) X chromosome has become inactivated [Fig. 1b]. We then picked single NPC colonies that can be clonally propagated and selected a clonal line (E6) in which the B6 X chromosome is inactivated. In this way, *Xist* expression levels from the inactive, B6 Xi can be increased by adding Dox to the culture media [Fig. 1b]. We also included two previously described NPC lines (CL30 and CL31) carrying an inactive B6 X chromosome^2^ [Extended Data Fig. 1e].

NPC clone E6 displays a high degree of escape, with 133 genes showing biallelic expression out of the 446 informative X-linked genes, whereas clones CL30 and CL31 show less escape, with 68 and 50 escapees, respectively [Extended Data Fig. 1e,f]. We checked whether the escapees identified in our NPC clonal lines have been previously shown to escape XCI [Supplementary Table 1; Extended Data Fig. 1g]. We thereby define three categories of genes: (i) ‘constitutive’ escapees (consistently escape XCI in all studies); (ii) ‘facultative’ escapees (variably escape XCI in different contexts) and (iii) ‘NPC-specific’ escapees (escape XCI only in NPC clones so far and behave like facultative escapees). Out of the 133 escapees identified in clone E6, 12 are constitutive escapees and 84 are facultative genes. All 12 constitutive escapees also escape in clones CL30 and CL31 in which we identified 38 and 28 facultative escapees, respectively [Extended Data Fig. 1h]. Only 10 NPC-specific genes were found to escape XCI in all clones; the majority escape variable in different clones [Extended Data Fig. 1g]. Characterising the extent and nature of escape variability across different clones, particularly in light of previous data, is very important to understand whether a specific subset of X-linked genes is sensitive to changes in Xist RNA levels regardless of the cell type or developmental context.

We went on to test whether Xist overexpression directly leads to silencing of these escapees in different NPC lines [Fig.1; Extended Data Fig. 1]. *Xist* RNA levels were increased on the Xi by adding Dox to the culture media for 3, 7, 14 and 21 days followed by RNA sequencing [Fig. 1c-f; Extended Data Fig. 2a-e]. Two to three replicates per time point were sequenced, and showed robust data reproducibility [Extended Data Fig. 2a-d]. *Xist* was efficiently upregulated upon Dox induction leading to a 6.9-7.5-fold enrichment in expression levels [Fig. 1d; Extended Data Fig. 2e]. Increased Xist RNA levels in Dox-treated NPCs were also observed by Xist RNA FISH analysis [Fig. 1c]. Upon *Xist* induction, the expression levels of escapees were found to be consistently reduced, demonstrating the capacity of Xist RNA to initiate escapee gene silencing in fully differentiated cells, even though this is not during the early developmental time window previously defined for Xist action [Fig.1e,f; Extended Data Fig. 2a-i]. After 3 days of Xist induction, 78 escapees out of 133 started to show silencing with a significant reduction in allelic ratio of at least 50% (binomial linear model, adjusted p < 0.05), while 89 genes were still expressed from the Xi (i.e. allelic ratio > 0.1). By day 14 and 21 of Dox treatment gene silencing became complete for up to 119 escapees with only 21 genes still showing residual escape (albeit at reduced level) after 21 days of Xist induction [Fig. 1f; Extended Data Fig. 2i].

Allele-specific differential expression analysis of X-linked genes in Dox-treated vs untreated NPCs showed significant downregulation of total levels of most escapees and confirms Xist-mediated silencing *in cis* only on the Xi, whilst the expression levels of homologous genes on the Xa remain unchanged [Extended Data Fig. 3a-b]. This observation suggests the lack of compensatory mechanisms that control the overall dosage of escapees, at least at the mRNA level [Extended Data Fig. 3a-c]. We also tested whether Xist upregulation affects the expression of autosomal genes. We found that only 3 genes were up-regulated and 13 down-regulated across the rest of the genome, showing that there are very limited effects of Xist upregulation on the expression levels of autosomal genes [Extended Data Fig. 3c].

Changes in allelic ratios of escapees over the Dox time course suggest different silencing dynamics for different categories of escapees [Fig 1.f,g, Extended Data Fig. 2f-h]. To better characterise the silencing dynamics of different escapees upon Xist overexpression in NPCs, we quantified the speed at which escape was lost upon Xist overexpression (silencing half-life); and asked whether after prolonged Dox treatment certain genes retained some expression (residual escape) [Fig. 1h; Extended Data Fig. 4a,b]. Briefly, we fitted two alternative exponential decay models, one in which we estimated the half-life *t1/2* of the allelic ratios approaching 0 (full silencing model) and one in which the decay approaches a residual escape level (residual escape model) [Fig. 1h]. For 31 genes (out of 122 with an exponential fit R2 > 0.3), the silencing kinetics were better explained by the model that includes a residual escape term, indicating that for a group of genes full silencing will not be established even after persistent Xist overexpression, while the majority become robustly silenced [Extended Data Fig. 4a]. For both groups, fully silenced and residual escape genes, we found that escapees showed a large variation in silencing speeds, with constitutive escapees being silenced more slowly than NPC-specific and facultative escapee genes, with the latter showing the widest range of silencing kinetics [Fig. 1i]. Similarly, residual escape was more frequent and higher amongst constitutive escapees than amongst facultative and NPC-specific escapees [Fig. 1k; Extended Data Fig. 4a,b].

Finally, we asked whether the expression levels of escapees or their position along the Xi account for the differences found in silencing upon Xist upregulation [Fig. 1j; Extended Data Fig. 4c,d]. We found no correlation between the general expression levels of escapees (Xa + Xi levels) and their efficiency of being silenced upon increased Xist expression (+Dox) [Extended Data Fig. 4c]. Similar results were obtained by taking into account only the Xi-specific expression levels of escapees [Extended Data Fig. 4d]. However, escapees pairs within 100 kb of each other showed more similar half-lives than randomly chosen pairs, implying that neighbouring escapees share similar responses to increased Xist levels [Fig. 1j]. We identified 11 regions with at least 3 escapees along the Xi characterised by different silencing speeds [Fig. 1e,f,j]. Interestingly, region 6 and 7, which include 12 and 13 genes respectively, correspond to neighbouring loci along the X chromosome, yet their dynamics of silencing is rather different, suggesting that proximity alone cannot fully explain the observed differences in silencing speed [Fig. 1e,j]. Altogether, we show that increased Xist RNA has the capacity to directly reduce the expression levels of escapees in post-XCI cells, leading to almost full loss of escape after 21 days of Dox induction in NPCs.

**Figure 1:**
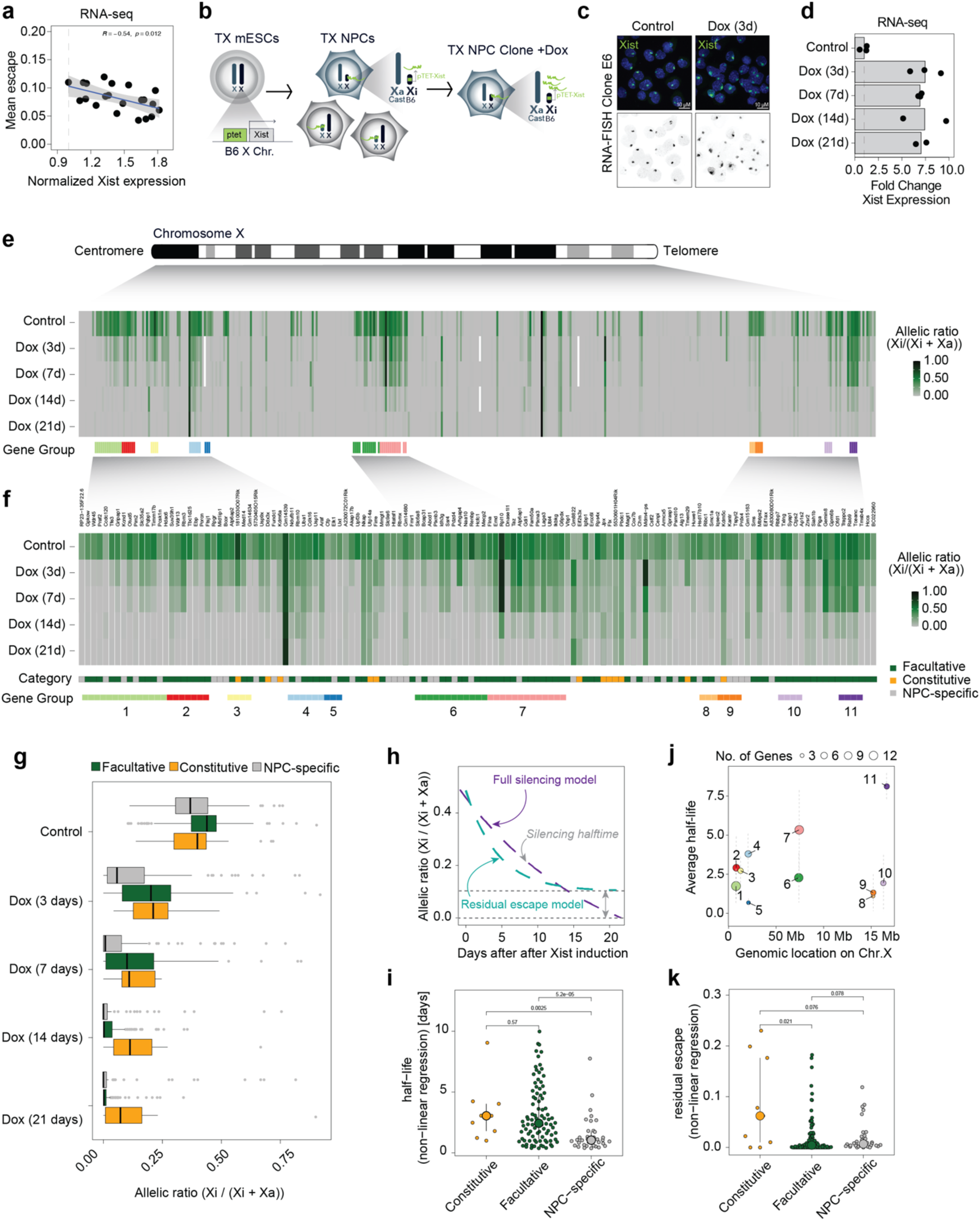
Increased levels of Xist RNA silences Xi escapees in NPCs. **a,** Scatterplot showing a correlation between average escape and *Xist* expression using RNA-seq data from 21 NPC clones (129/Sv x Cast/Eij genetic background). Mean escape is calculated as the average allelic ratio (Xi/(Xi+Xa)) across 439 informative genes. Normalised *Xist* expression is calculated as library-size scaled counts per million (CPM), divided by the value for the lowest clone. *R* specifies Pearson’s correlation coefficient, the p-value is given by a correlation test. **b,** Experimental outline: TX ESCs carrying a ptet tetracycline-responsive promoter upstream of the Xist gene on the B6 X chromosome were differentiated NPCs without doxycycline. Single clones carrying the inactivated B6 allele were picked and expanded and Xist RNA levels were increased by adding doxycycline to the culture media. **c,** FISH for Xist RNA (green) in NPC clone E6 in untreated conditions (Control) and after 3 days of doxycycline treatment (Dox 3d). DNA is stained with DAPI. **d,** RNA-seq data showing the fold change in Xist expression (normalised CPM) compared to untreated cells across the time course of doxycycline treatment. Data relative to the mean of measurements in clone E6 is shown. **e,** Schematic of the mouse X chromosome and heatmap showing X-linked transcript allelic ratios in untreated clone E6 and after 3, 7, 14 and 21 days of doxycycline treatment. Allelic ratio indicates the fraction of reads from the Xi (ratio=1: Xi monoallelic expression; ratio=0: Xa monoallelic expression; ratio=0.5: biallelic expression; ratio > 0.1: escape). Gene groups are defined as contiguous groups of escapees within 100 kb of each other (see Methods). **f,** Heatmap showing the allelic ratio of 134 escapees identified in clone E6 and shown in **e**. Escapees are assigned to three different categories as indicated in Extended data Fig.1g and described in Methods). The escape category for each gene is indicated below the heatmap together with the zoom in of the gene groups shown in **e**. **g,** Box plot showing the changes in allelic ratios for different escape categories across the time course of doxycycline treatment. Data of clone E6 is shown. **h,** Schematic of the exponential decay models used to study gene silencing kinetics. Data can be described by a full silencing model (blue, allelic ratio approaches 0) or a residual escape model (green allelic ratio approaches value > 0.1. The steepness of the curve corresponds to the genes silencing half-life). **i,** Beeswarm plot showing the distributions of silencing half-life fit to all escapees using the offset model and stratified by escapee classification. P-values are computed using Wilcoxon’s Rank Sum test (not adjusted for multiple testing). **j,** Silencing Half-lives per gene group as shown in (**f**). Large dots show mean and whiskers standard deviations across genes in the group. **k,** Beeswarm plot showing the distributions of residual escape parameters fit to all escapees using the offset model and stratified by escapee classification. P-values are computed using Wilcoxon’s Rank Sum test (not adjusted for multiple testing).

### Xist-mediated silencing in NPCs is SPEN-dependent

We asked whether the mechanisms underlying Xist-mediated silencing in NPCs are the same as those that control XCI initiation at the onset of the process in mESCs and during early development when the transcriptional repressor SPEN plays an essential role ^2^. Indeed, SPEN knockout embryos fail to initiate iXCI and phenocopy mutants embryos lacking *Xist* ^2,38^. To address whether Xist-mediated escapee silencing in post-XCI cells is SPEN dependent, we used two NPCs clones (CL30 and CL31) in which both endogenous *Spen* alleles are tagged with the AID degron domain. This allows SPEN acute degradation upon addition of the plant hormone auxin (indole-3-acetic acid; IAA) to the culture media ^2^. SPEN was depleted in the two NPC clones for 2 days before inducing *Xist* upregulation for 3 and 7 days [Fig. 2a]. Depletion of SPEN in absence of Xist upregulation led to moderate upregulation of escapees, confirming that in female NPCs SPEN dampens their expression levels on the Xi, as we previously observed [Extended Data Fig. 5a; Dossin et al., 2020]. *Xist* was efficiently upregulated upon combined Dox and Auxin treatment, reaching up to 10-fold enrichment compared to untreated NPCs [Extended Data Fig. 5b]. However, in absence of SPEN, the higher levels of Xist RNA did not alter the expression levels of escapees, contrary to the Xist-mediated silencing seen in the presence of SPEN [Fig. 2b, Extended Data Fig. 5a]. Notably, all three categories of escapees were unaffected by Xist in the context of SPEN depletion, suggesting that the capacity of Xist RNA to modulate the expression levels of all escapees in NPCs is strictly SPEN-dependent [Fig. 2c,d]. Our data thus reveal that SPEN is essential for Xist-mediated silencing of escapees in differentiated cells, similarly to the situation during early embryonic development which is when XCI is initially established.

### Xist-mediated silencing of escapees leads to loss of TAD-like domains on the Xi

Our previous studies revealed that clusters of facultative escapees reside in TAD-like structures on the inactive X chromosome, which is otherwise devoid of TADs in differentiated cells ^57,61^. However, the relationship between Xist RNA coating, gene silencing, and 3D topology on the inactive X chromosome is not yet fully understood. In particular, whether the topological organisation of escaping clusters is a cause or a consequence of escapee expression, remains an open question.

Here we addressed how chromosome topology responds to changes in transcriptional activity of escapees, upon increased *Xist* expression with or without SPEN depletion. We performed allele-specific Capture Hi-C analysis of >3 Mb genomic regions encompassing the TAD-like domains previously observed on the Xi in clonal NPCs ^57^ [Fig. 3a,b]. One of these (the ‘*Mecp2-Hcfc1* cluster’) spans *∼*800 kb and includes 19 facultative and 6 NPC-specific escapees [Fig. 1e,f]. In clone E6, all 25 genes escape in this region, whereas in CL30.7 only 10 genes are biallelically expressed [Extended Data Fig. 1e]. The second, *’Kdm5c* cluster’, spans *∼*500 kb and includes the constitutive escapee *Kdm5c* together with 12 facultative escapees [Fig. 1e,f]. As expected, we observed a general lack of TADs at silenced genes on the Xi, whereas clusters of escaping genes were organised into 3D TAD-like structures that resemble the topological organisation of the Xa [Fig. 3a,b]. After 7 days of Dox induction, the 3D topology of the Xa at the *Mecp2-Hcfc1* cluster remains unchanged [Extended Data Fig. 6a] whereas TAD-like structures at the same locus on the Xi become attenuated, but not completely lost [Fig. 3a]. In particular, escapees that are robustly silenced show more pronounced loss of topological organisation whereas genes that are only dampened but remain transcriptionally active retain their structural organisation [Fig. 3a]. Interestingly, the first group of escapees lies in region 6, characterised by relatively faster gene silencing than neighbouring region 7 which includes escapees showing longer silencing half-life and retaining topological organisation after 7 days of Dox treatment [Fig. 1j; Fig. 3a]. However, by day 21of Xist upregulation most escapees in the *Mecp2-Hcfc1* cluster are effectively silenced and TAD-like structures are lost across the entire region [Fig. 3a].

**Figure 2:**
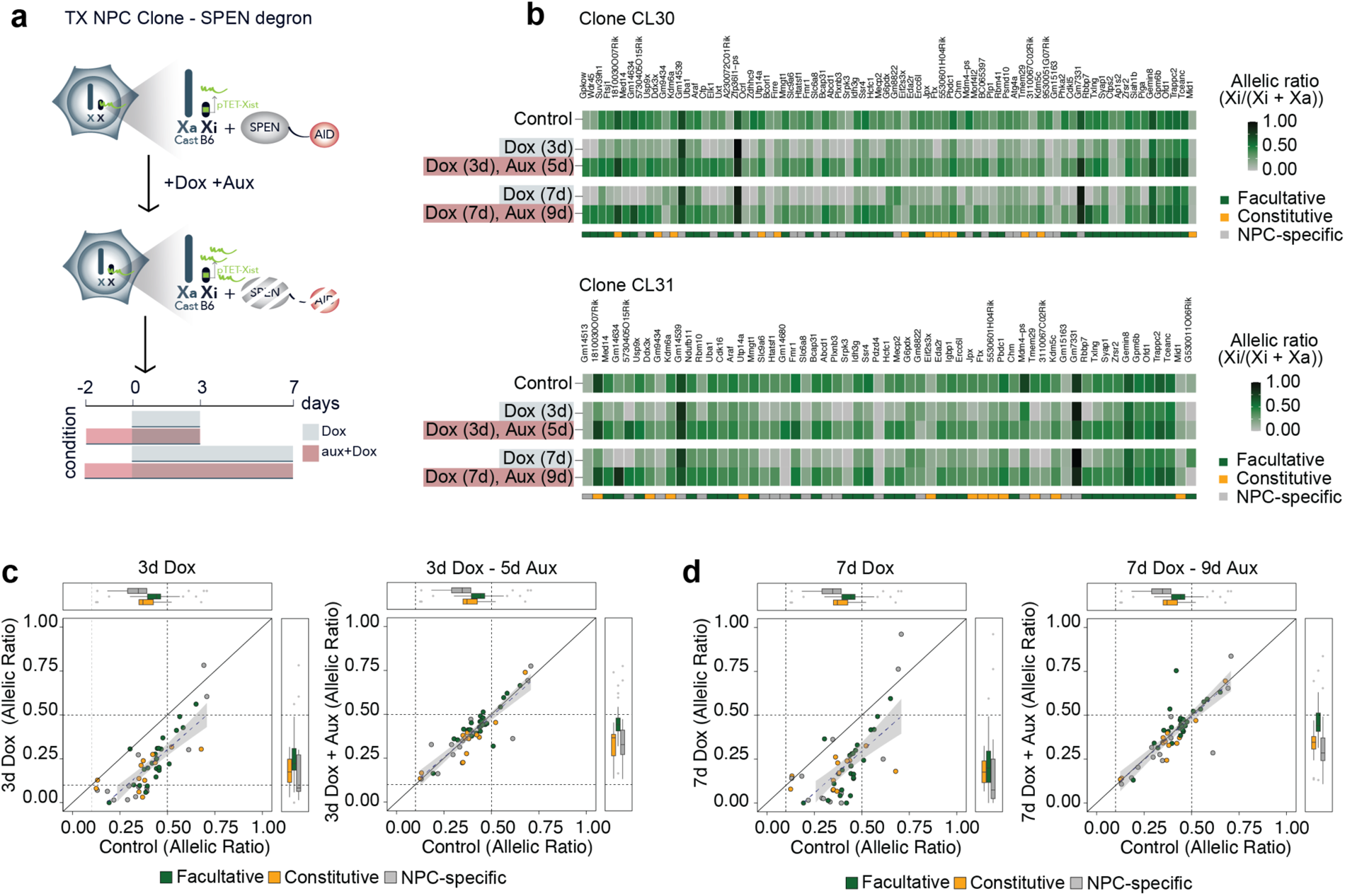
Xist-mediated silencing in NPCs is SPEN-dependent. **a**, Experimental outline: Xist RNA levels were increased in NPC clones carrying a SPEN-AID degron (Dossin at al. 2020). SPEN was depleted by adding auxin (aux) to the culture media for 2 days before inducing Xist upregulation with doxycycline (Dox) for 3 or 7 days in the presence of auxin. **b,** Heatmap showing allelic ratios of escapees upon Xist induction for 3 days (Dox (3d)) and 7 days (Dox (7d)) and in combination with auxin treatment (Dox (3d), Aux (5d) and Dox (7d), Aux (9d)). Data from NPC clones CL30.7 and CL31.16 is shown. The escape category for each gene is indicated below the heatmap. **c-d,** Dot plots representing changes in allelic ratios upon doxycycline and auxin treatments for the 3 day (c) and 7 day timepoints (d). Shown are the average allelic ratios of genes escaping in both CL30.7 and CL31.16 clones. The diagonal line represents no change compared to an untreated cell line.

We observed similar results at the *Kdm5c* cluster [Fig. 3b, Extended Data Fig. 6b]. Here, 7 days of Xist upregulation resulted in efficient silencing of all 11 facultative and NPC-specific escapees in the cluster, although *Kdm5c* which is a constitutive escapee resists complete inactivation [Fig. 3b]. Similarly to the *Mecp2-Hcfc1* cluster, upon Xist upregulation the TAD-like structures encompassing all escapees at this locus are lost. Only a long-range looping interaction upstream of *Kdm5c* is maintained, probably allowing this locus to engage in long-range contacts with other escapee loci, as previously suggested [Fig. 3b] ^34,57^. These results suggest that the local 3D structures at escapee clusters on the Xi are the result of ongoing transcription at these loci.

Xist RNA has previously been proposed to play a role in shaping the conformation of the Xi, independently of transcription at least in part ^15,34^. To assess whether the effect we saw on X-chromosome structure was due to increased Xist RNA levels or to gene silencing we analysed cells with or without SPEN. We performed allele-specific Capture Hi-C experiments in NPC clone CL30 containing the SPEN-AID degron [Fig. 3c]. No differences in 3D topology on the Xa were observed upon doxycycline (Xist up-regulation) and auxin (SPEN depletion) treatment [Extended Data Fig. 6c]. Inducing Xist upregulation for 21 days while triggering the acute depletion of SPEN results in no loss in local 3D topology of the Xi. Escapee genes within the *Mecp2-Hcfc1* cluster remain actively transcribed and organised in 3D TAD-like domains, even though Xist RNA levels are 10-fold higher in Dox-treated cells [Fig. 3c].

Altogether, our results reveal that increased Xist RNA levels only lead to loss of TAD-like structures on the inactive X, in the presence of SPEN as Xist RNA upregulation in the absence of SPEN does not affect the 3D topology of these loci. This demonstrates that Xist RNA is capable of shaping the conformation of the Xi at escapee loci, only in the context of transcriptional repression of X-linked genes.

### Silencing of most escapees becomes Xist-independent after prolonged Xist upregulation

Another key question we wished to address was whether Xist-mediated silencing of escapees in NPCs is reversible and strictly dependent on persistent Xist overexpression, or whether silencing eventually becomes irreversible and epigenetically locked in. To address this, we induced *Xist* upregulation for 3, 7, 14, and 21 days in NPC clone E6 followed by 7 days of washout and we performed RNA sequencing [Fig. 4a-d]. During the time course NPCs continued to actively divide and we did not notice any obvious effect on cell proliferation. After Dox washout Xist expression levels return to the levels found in untreated cells prior to Dox-induced overexpression [Fig. 4b,c]. Our analysis reveals progressive irreversibility of silencing upon longer periods of Xist upregulation, with different escape categories following different dynamics of reactivation after Dox wash-out [Fig. 4d-f]. After 7 days of Xist upregulation followed by 7 days of Dox washout, 95 escapees out of 133 show reversible silencing and become reactivated (i.e. allelic ratio >0.1 and at least 50% of original escape level), whereas 16 genes are only partially irreversible (i.e. <50% of original escape level) and 22 escapees are irreversibly silenced [Fig. 4f]. Amongst the irreversible genes, 13 genes are NPC-specific and 9 facultative genes [Fig. 4f]. Notably, all 12 constitutive escapees are fully reversible at this time point [Fig. 4f]. After 14 days of Xist upregulation, the repressed state of 57 escapees becomes irreversible when checking after 7 days of washout [Fig. 4f]. Of these genes, 25 are NPC-specific, 30 facultative and 2 constitutive [Fig. 4f]. The number of escapees becoming irreversibly silenced remains mostly unchanged, across all categories after 21 days of Xist induction (compared to 14 days), suggesting that for irreversibly-silenced genes, Xist RNA becomes dispensable after 2 weeks of upregulation [Fig. 4f]. Notably, most constitutive escapees are still fully reactivated upon washout following 21 days of Xist upregulation. Thus, although constitutive escapees are sensitive to Xist RNA levels, they remain resistant to complete XCI [Fig. 4df].

We also investigated whether the observed differences in reactivation across different escapees reflect the differences in silencing dynamics observed in the 21 days Dox time course [Fig. 4g]. We found that escapees that undergo fast silencing upon increased Xist levels are less prone to reactivate, whereas slowly silenced genes tend to retain the capacity to become reactivated, when Xist returns to its basal expression levels [Fig. 4g].

**Figure 3:**
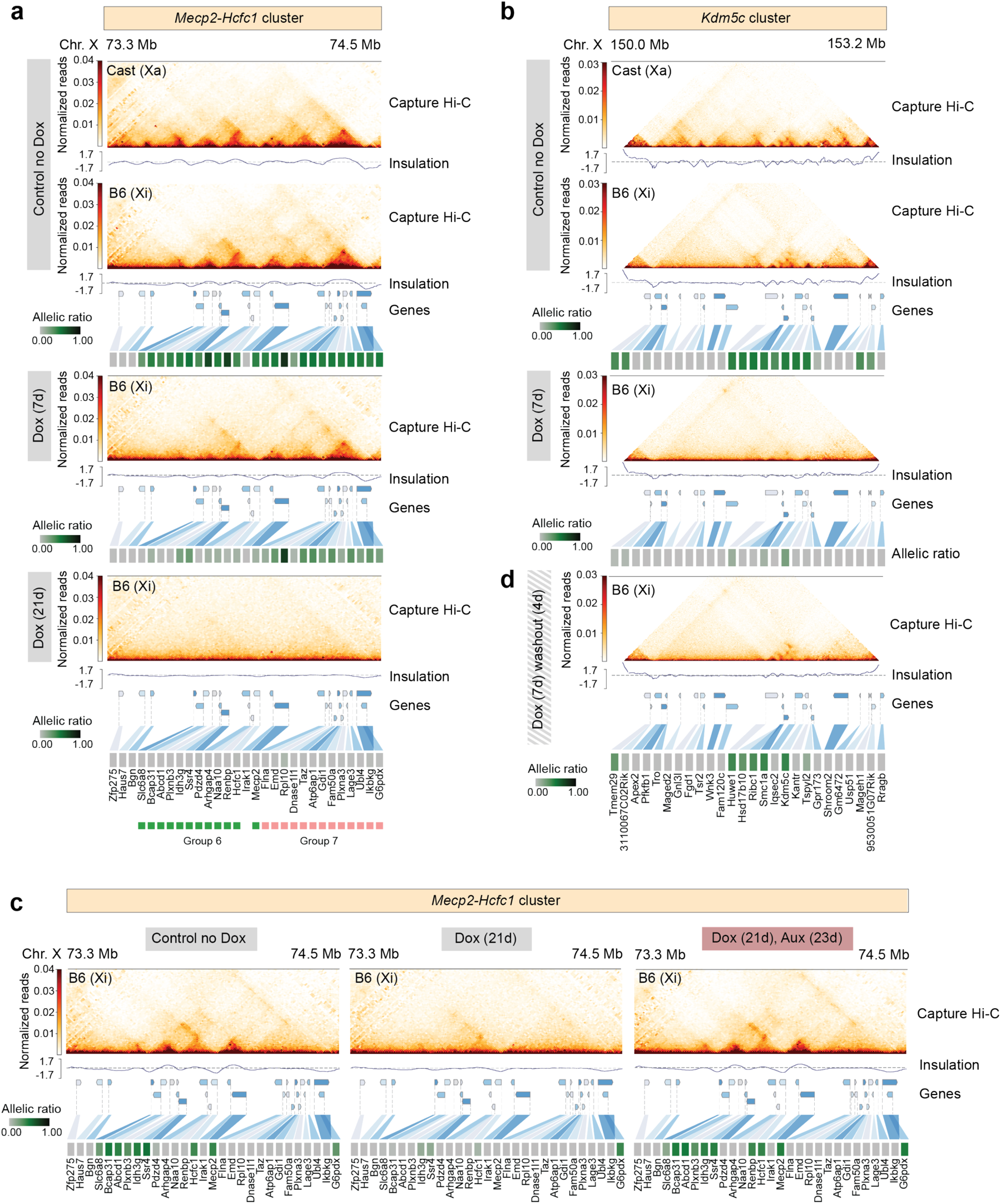
Xist-mediated silencing of escapees leads to loss of TAD-like domains on the Xi. **a-b**, Capture Hi-C interactions and insulation score at the **(a)** *Mecp2-Hcfc1* and **(b)** *Kdm5c* cluster in clone E6 prior to and upon doxycycline treatment. Capture Hi-C interactions are shown for the active (Xa) and the inactive (Xi) X chromosomes in untreated conditions (control no Dox) (top) and for the Xi upon doxycycline treatment for 7 days (middle) and 21 days (bottom) for the *Mecp2-Hcfc1* cluster and 7 days for the *Kdm5c* cluster. Capture Hi-C data is shown at 10-kb resolution. Heatmaps showing allelic ratios for 29 **(a)** and 27 **(b)** X-linked genes included in the captured regions are shown. Escapees belonging to groups 6 (green) and 7 (pink) within *Mecp2-Hcfc1* cluster are highlighted. **c,** Capture Hi-C interactions and insulation score at the *Mecp2-Hcfc1* cluster for Xi in clone CL30.7. Left to right: untreated cells (control no Dox); 21 days of doxycycline induction; 23 days of auxin treatment in combination with 21 days of doxycycline induction. Capture Hi-C data is shown at 10-kb resolution. Heatmaps showing allelic ratios for 29 X-linked genes included in the captured region are shown. **d,** Capture Hi-C interactions and insulation score at the *Kdm5c* cluster for Xi in clone E6 after 7 days of doxycycline treatment followed by 4 days of doxycycline washout. The 27 X-linked genes included in the captured regions are shown.

The latter situation is illustrated by the 4 escapees included in group 11, which show the longest silencing half-life and get fully reactivated after 21 days of Dox treatment [Fig. 4g]. On the other hand, gene silencing of escapees in regions 5, 8, and 9 become irreversible after 21 days of Xist upregulation [Fig. 4g]. As we observed for escapee silencing dynamics, the genes in regions 6 and 7 also respond differently to Dox washout, despite being in close proximity to each other on the Xi [Fig. 4g]. Given that gene silencing is largely reversible after 7 days of Xist upregulation at both *Hcfc1-Mecp2* and *Kdm5c* clusters organised in TAD-like structures on the Xi [Fig. 3a,b Fig. 4d], we tested whether reducing Xist to basal levels (Dox washout) would restore their topological organisation. Indeed, the 3D spatial organisation of these loci that is either attenuated or lost after 7 days of Xist upregulation [Fig. 3a,b] becomes re-established when active transcription is restored on the Xi upon Dox washout [Fig. 3d, Extended Data Fig. 6d], whereas the 3D topology of the same loci on the Xa remains unchanged throughout the experiment [Extended Data Fig. 6b]. These data reinforce the observation that the emergence to TAD-like structures at escaping loci is the direct consequence of active gene transcription.

In summary, we showed that in NPCs, silencing of most escapees become irreversible and Xist-independent after 14 days of Xist upregulation, and that constitutive escapees are mostly reversibly silenced by Xist upregulation whilst the other categories of escapees show more variable responses in terms of gene reactivation after Dox washout.

### Irreversible silencing of escapees is facilitated by increased promoter methylation

Given the known role of DNA methylation in stably locking in the silent state of genes on the Xi ^12^, we assessed whether irreversible silencing of escapees is accompanied by DNA methylation. On the Xi, promoters CpG islands (GCI) of silent genes - but not of escapees - are hypermethylated, whereas CpG methylation of gene bodies is found at transcribed escapees on both Xa and Xi chromosomes, similar to other active genes in the genome ^63,64^.

We assessed whether increased levels of Xist RNA lead to changes in DNA methylation at escapees and whether this epigenetic feature can account for the transition from reversible to irreversible gene silencing upon Xist overexpression in NPCs. We performed allele-specific DNA-methylation analysis by Next Enzymatic Methyl-seq (EM-seq), after 7 and 21 days of Xist upregulation and Dox washout [Fig. 5a,b]. We found a progressive increase in DNA methylation at CpG-dense regions chromosome-wide upon Xist overexpression, suggesting that prolonged high levels of Xist RNA leads to *de novo* DNA methylation in NPCs [Fig. 5a]. We also found a significant decrease in DNA methylation at escapee gene bodies on the Xi (compared to silent genes on the Xi, where no change was seen), indicating that as escapees become repressed they acquire the epigenetic features of silenced genes on the Xi (i.e. hypermethylated CpG islands and hypomethylated gene bodies)[Fig. 5a,b]. These changes in DNA methylation at CpG islands and gene bodies of escapees were found to remain unchanged when Xist RNA levels were reduced to basal levels upon Dox washout for 7 days [Fig. 5a,b].

To gain DNA methylation information at a higher resolution and assess the methylation status of CpG islands at promoters of escapees that are either reversibly or irreversibly silenced, we also performed targeted capture methylation sequencing (EM-Cap) ^65^ [Fig. 5c,d]. We focused on the *Mecp2-Hcfc1* cluster as it includes escapees that differentially respond to varying Xist RNA levels. To assign CpG islands to promoters of escapees on the Xi we used allele-specific ATAC-seq peaks obtained from clone E6 before inducing Xist upregulation [Fig. 5c,d]. In this way, we were able to observe higher levels of DNA methylation at CpG-dense promoters of escapees that are irreversibly silenced compared to partially irreversible genes after 21 days of doxycycline treatment [Fig. 5c,d]. DNA methylation levels remain stable after doxycycline washout [Fig. 5c,d].

**Figure 4:**
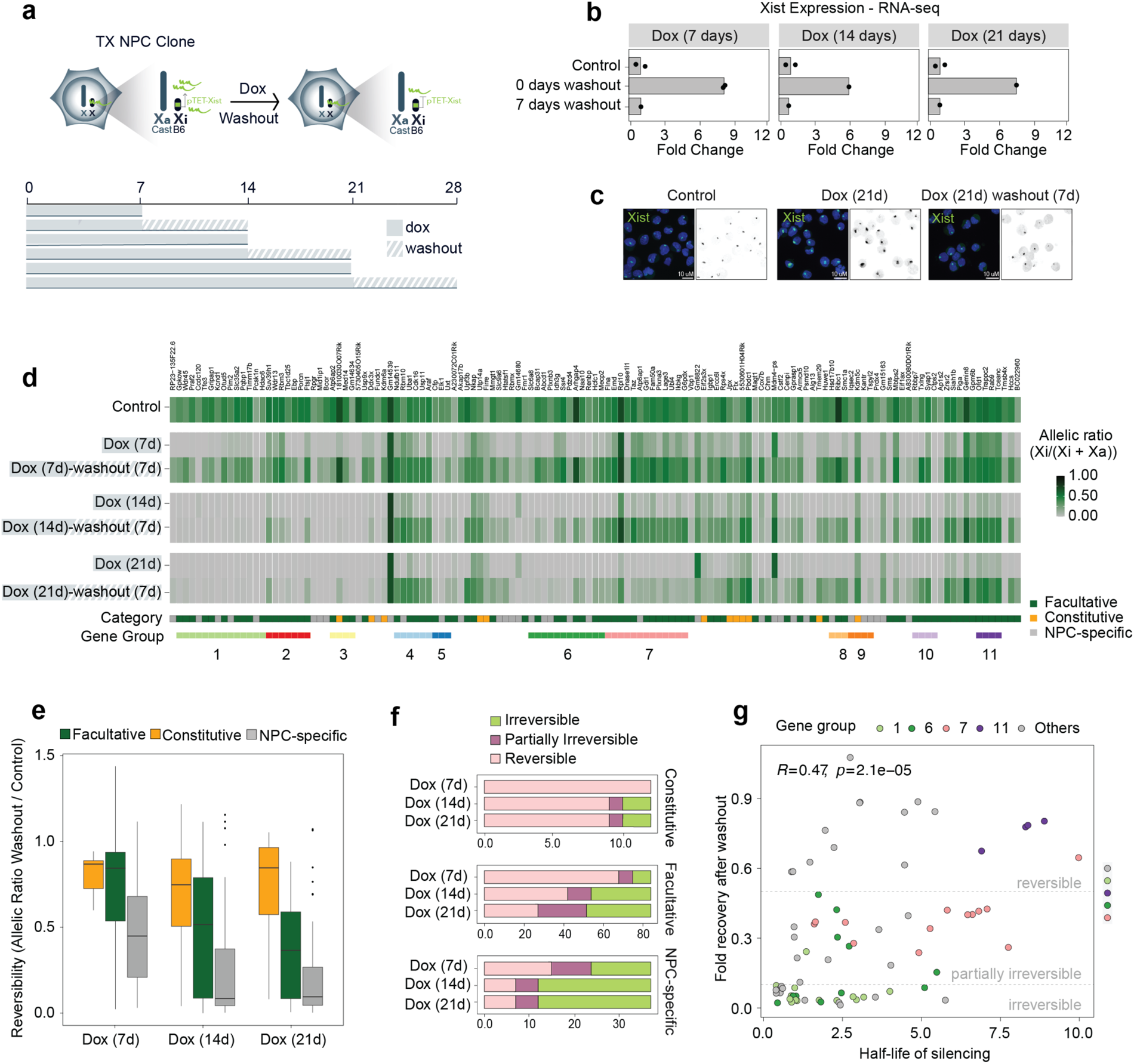
Silencing of most escapees becomes Xist-independent after prolonged Xist upregulation. **a**, Experimental outline: Xist upregulation was induced by doxycycline treatment for 7, 14, and 21 days following 7 days of doxycycline washout. **b,** RNA-seq data showing fold changes in Xist expression (normalised CPM) compared to untreated control cells after 7, 14 and 21 days of doxycycline treatment followed by 7 days of doxycycline washout. Data relative to the mean of measurements in clone E6 is shown. **c,** FISH for Xist RNA (green) in NPC clone E6 in untreated conditions (Control), after 21 days of doxycycline treatment (Dox 21d), and following 7 days of doxycycline washout (Dox 21d - washout 7d). DNA is stained with DAPI. **d,** Heatmap showing allelic ratios of 133 escapees identified in clone E6 upon doxycycline treatment for 7, 14, and 21 days and following 7 days of doxycycline washout after these time points. Escapees are assigned to three different categories as described in Supplementary Figure 1 (see Methods). **e,** Box plots quantifying the reversibility of escape per gene for each time point, by computing the ratio of allelic ratios after washout divided by the allelic ratio in untreated samples. **f,** Boxplots showing a classification of genes into irreversible, partially irreversible and fully reversible escapees across the silencing time course. Genes are considered reversible when they reach at least 50% of untreated escape as well as an allelic ratio > 0.1 after washout. Similarly, they are considered partially irreversible when they reach 10%-50% of untreated escape and an allelic ratio > 0.1, and irreversible otherwise. **g,** Scatterplot comparing silencing half-lives (see Fig. 1) to fractional reversibility (e). Genes in a subset of local gene groups are highlighted to show their coordinated behaviour.

In summary, we show that when increased for at least 14 days Xist RNA levels lead to irreversible silencing of escapees, and this is accompanied by loss and gain of DNA methylation at gene bodies and promoters, respectively. These changes mirror the patterns observed at genes that are robustly and stably silenced during XCI. On the contrary, escapees that remain partially irreversible upon Xist upregulation show a smaller gain of DNA methylation at their promoters and this is reflected by their capacity to become fully re-expressed when basal Xist RNA levels are restored even after 3 weeks of Xist induction.

**Figure 5:**
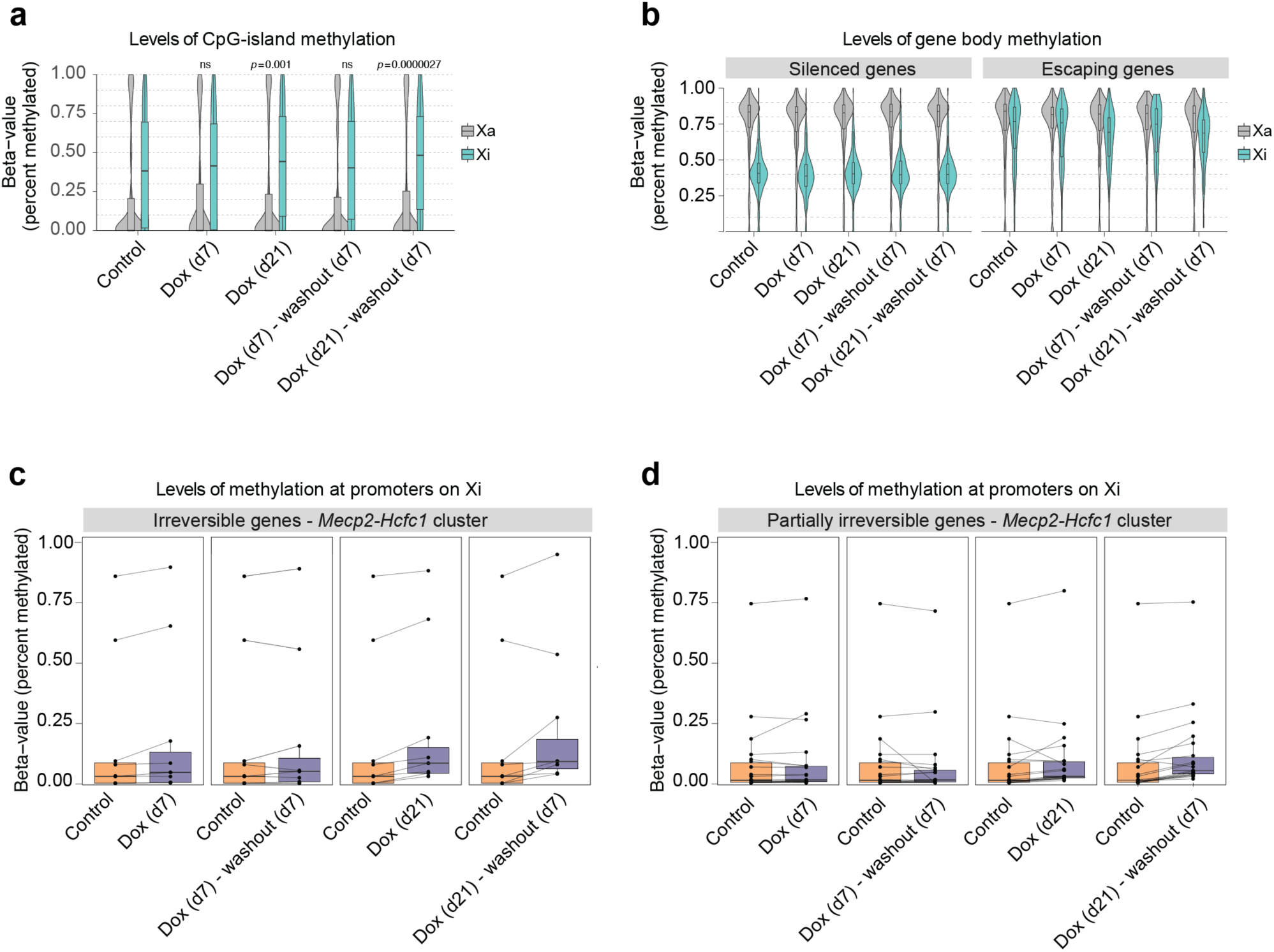
Irreversible silencing of escapees is maintained by DNA methylation. **a**, Box- and Violin plots showing allele-specific DNA methylation levels of CpG islands for control, doxycycline-treated and washout samples. For each GpG island, the average fraction of methylated cytosines is calculated across CpG sites from haplotype-informative reads. Shown are active (Xa=Cast) and inactive (Xi=B6) chromosomes separately. P-values are calculated using a paired Wilcoxon’s rank sum test. **b,** As (a), but showing methylation levels across gene bodies for silenced and escaping genes, as derived from the gene expression data in Fig. 1. **c,** Quantification of DNA methylation on promoters of irreversible escapees in the *Mecp2/Hcfc1*-cluster. Promoters are defined as regions of accessible chromatin as measured by ATAC-Seq that overlap (-500bp, 1000bp)-intervals around annotated TSSs. DNA methylation is defined as the average fraction of methylated Cs across the ATAC-peak region. Data is shown paired to untreated cells**. d,** As (c) for genes that are partially irreversible.

### Xist levels modulate X-linked dosage *in vivo*

The above studies were all performed on NPC cell lines *in vitro*. In order to assess whether Xist RNA levels can similarly modulate the expression levels of escapees *in vivo*, we investigated the impact of increasing Xist expression during preimplantation mouse development. To this end, we used mice that carry a doxycycline-inducible endogenous Xist allele (TX) and the tetracycline-responsive transactivator (rtTA) integrated at the ubiquitously expressed *Rosa26* locus ^60^. These mice were originally used to establish the TX1072 mESCs ^59^ and the NPC derivatives used in this study. We crossed TX/Y males heterozygotes for the rtTA transactivator (X^ptet^/Y; R26^rtTA/WT^) with wild type *Mus musculus molossinus* (JF1) females to obtain F1 hybrid embryos, in order to distinguish maternal and paternal alleles [Fig. 6a]. As iXCI always leads to the inactivation of the paternal X chromosome (Xp), this strategy allows us to perform whole embryo RNA-seq [Fig. 6a].

We collected F1 embryos at E2.5 and E3.5 stages during pre-implantation development and cultured them *in vitro* for 24 hours in the presence of Dox before performing RNA-seq at E3.5 and E4.5, respectively [Fig. 6a]. Given the genotype of TX/Y males, all F1 female embryos inherited the dox-inducible Xist allele but Xist upregulation was induced only in embryos carrying the rtTA transactivator which is essential for the inducible system to work [Fig. 6a]. This system provided us with control embryos in which Xist levels remained unchanged upon doxycycline induction [Fig. 6a]. We selected E3.5 and E4.5 developmental stages to avoid the establishment phase of iXCI, which starts at the 4-cell stage, in order to assess the impact that increased Xist RNA levels have, once iXCI has already occurred [Fig. 6a] ^38^. Indeed iXCI is largely complete by the early blastocyst state (E3.5) with most genes being partially or fully silenced, and some genes escaping XCI ^23,38^.

Increased levels of Xist RNA in female embryos carrying rtTA was detectable by RNA FISH as larger and more intense clouds, similarly to what we observed in doxycycline-treated NPCs [Fig. 6b,c]. Xist upregulation was also observed by whole embryo RNA-seq at E4.5, although we noticed a degree of variability in Xist RNA levels between different embryos and in particular in rtTA negative controls at E3.5 [Fig. 6d]. Principal components analysis (PCA) of X-linked allelic ratios across all sequenced embryos clustered them according to their developmental stage [Fig. 6e]. Notably, rtTA positive embryos show lower X-linked allelic ratios when compared to their stage-matching controls at E3.5 and E4.5, suggesting that increased Xist RNA levels lead to higher levels of silencing of X-linked genes that are expressed from the Xi at these stages of iXCI *in vivo* [Fig. 6f].

To address the impact of Xist upregulation on the expression levels of escapees on the Xi, we focused on E4.5 embryos as XCI is complete at this stage, with 203 genes being inactivated out of 343 informative genes compared with 133 inactivated genes at E3.5 [Fig. 6g] (median allelic ratio >0.1). We detected 135 escapees at E4.5 (i.e. allelic ratio >0.1 in at least 50% of embryos) [Fig. 6g,h]. These genes include 9 constitutive escapees and 87 facultative escapees [Fig. 6h]. Eight of the constitutive escapees identified in embryos and 61 of the facultative genes were also detected in NPCs, respectively [Fig 6h]. Out of the 31 NPC-specific escapees identified in NPCs, 13 escaped XCI in embryos as well, confirming their facultative nature. We also identified 27 embryo-specific genes that were reported as inactivated genes in our reference list [Supplementary Table 1]. Upon Xist upregulation most escapees become downregulated on the Xi, with 117 genes out of 135 showing decreased expression levels [Fig. 6h]. Similarly to what we observed in NPCs, different categories of escapees showed different responses to increased Xist RNA levels [Fig. 6h,i]. Constitutive escapees are less sensitive to Xist upregulation once again highlighting their intrinsic capacity to evade Xist-mediated silencing in different contexts [Fig. 6i,j]. Interestingly, embryo-specific escapees show variable responses to Xist upregulation with the majority of genes showing a reduction in allelic ratio but also a subset of them being mostly unaffected [Fig. 6i,j]. A closer examination of these latter genes revealed that two of these them, *Rho5x* and *Fthl17f* (also known as *Gm5635*), are imprinted genes only expressed from the Xp ^66,67^ and have previously been seen to be expressed on the Xp at this stage of development ^38^ [Fig.6j]. Finally, as observed in NPCs, facultative escapees (including NPC-specific escapees) are the most efficiently silenced genes upon Xist upregulation in E4.5 embryos [Fig. 6i, j].

Thus, these experiments confirm our *in vitro* findings and prove that Xist RNA levels can control the degree of escape from XCI *in vivo* for the majority of escapees although constitutive escapees are affected to a lesser extent.

**Figure 6:**
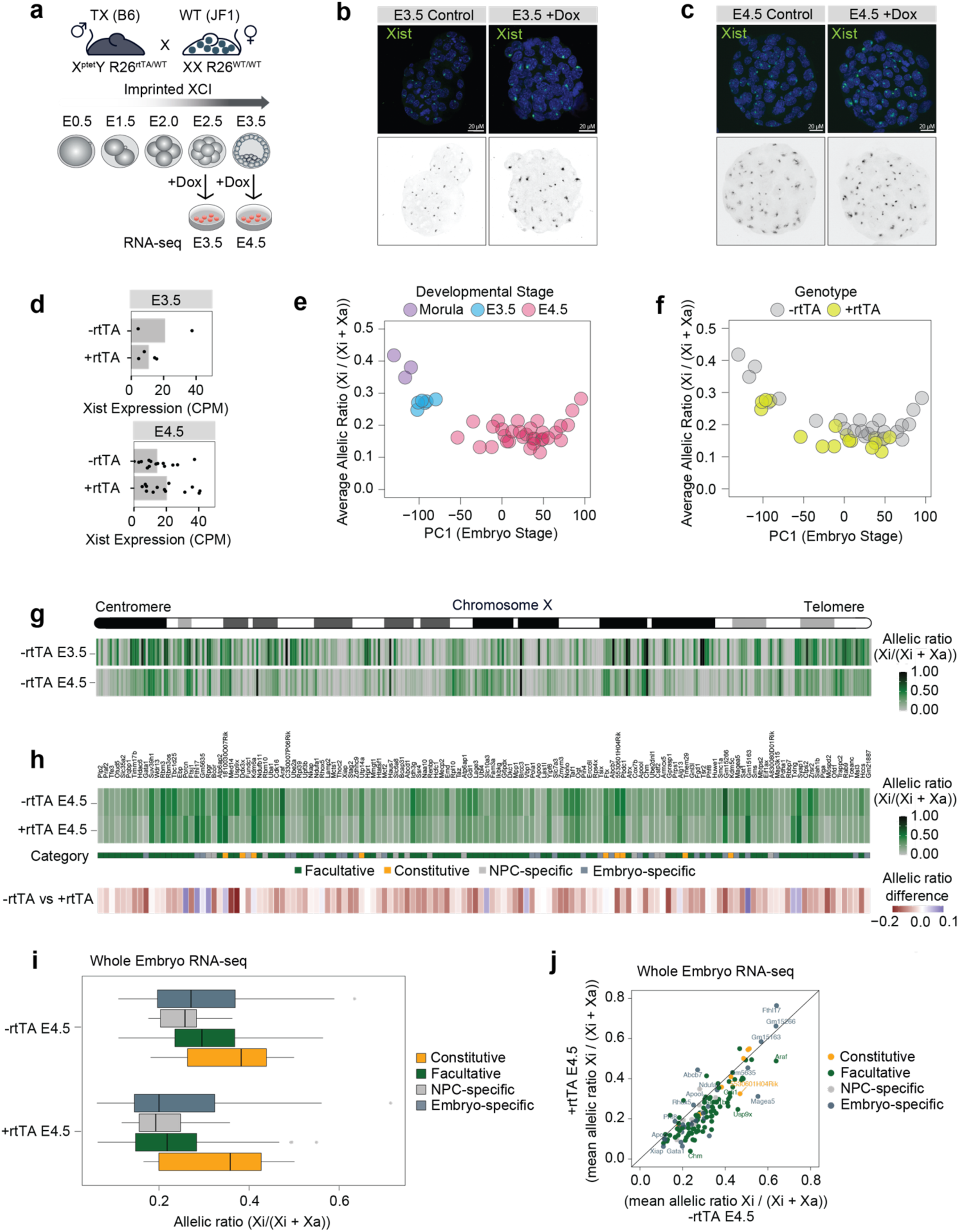
Xist levels modulate X-linked dosage *in vivo*. **a**, Experimental outline: males TX B6 mice (X^ptet^Y; R26^rtTA/WT^) were crossed with WT JF1 females. F1 embryos were collected at E2.5 and E3.5 and cultured for 24 hours while adding doxycycline to the culture media. RNA-seq was performed at E3.5 and E4.5. **b-c,** FISH for Xist RNA (green) in (**b**) E3.5 and (**c**) E4.5 XX embryos obtained by crossing males TX B6 mice (X^ptet^Y; R26^rtTA/rtTA^) with WT JF1 females. DNA is stained with DAPI. **d,** Whole embryo RNA-seq at E3.5 and E4.5 showing Xist RNA levels (normalized CPM) in females carrying the rtTA transactivator compared to WT females (Methods). Each dot represents one embryo. **e,** Scatterplot showing average escape across the embryo stage as defined by principal component analysis on the whole transcriptome (Methods). Average escape is defined as the mean allelic ratio of all expressed X-linked genes. Developmental stage annotations are defined based on embryo morphology. **f,** As (e), but showing rtTA genotypes based on RNA-seq. Embryos are classified as +rtTA by PCR and if at least 50 reads mapped to the rtTA transgene (Methods). **g,** Schematic of the mouse X chromosome and heatmap showing the allelic ratios of X-linked genes expressed in female embryos at E3.5 and E4.5. For each gene and developmental stage, the median allelic ratio is shown. **h,** Heatmap showing the mean allelic ratios of 135 escapees in E4.5 embryos carrying rtTA (+rtTA) and in matching controls (+rtTA). Escapees are categorised as in Extended data Fig.1g, with genes additionally called “embryo-specific” if they show an allelic ratio > 0.1 in >50% of -rtTA embryos and do not escape in NPCs. Below, the mean difference in allelic ratios between -rtTA and +rtTA embryos are shown. **i,** Box plots showing mean allelic ratios for -rtTA and +rtTA embryos for the different escapee categories. **j,** Scatterplot showing the same data as in (i), but directly comparing allelic ratios in -rtTA and +rtTA embryos.

## DISCUSSION

Our study demonstrates that Xist RNA levels can directly modulate expression levels of all X-linked escapees in differentiated cells, via Xist’s key silencing partner, SPEN, and beyond the developmental time window previously assumed to be Xist’s main time of action. These findings pave the way to understanding how natural fluctuations in Xist expression levels, that may arise during development, in adult tissues or pathological situations such as in cancer, might impact X-linked escapee gene expression and protein dosage.

Recent work in humans and mice showed that deletion or knock down of Xist/XIST, for example in human mammary stem cells^48^, or in the murine hematopoietic system^53^, can lead to epigenetic changes that have small but significant effects on X-linked gene expression. Even a small increase in X-linked gene expression can result in important phenotypic consequences, for example on mammary cell differentiation potential, leading to female-specific cancer development ^48,53^. This is illustrated by the recent example of MED14, a facultative Xi escapee in both mouse and humans ^25,68^, the up-regulation of which from the Xi results in increased levels of Mediator complex with major consequences on mammary stem cell differentiation and breast cancer^48^. Similarly, the re-activation of a subset of X-linked escapees leads to female-specific leukaemia in mice^53^.

We now provide direct evidence that the levels of Xist RNA participate in the regulation of X-linked escapees in a post-developmental context, potentially throughout life. This implies that natural genetic variation at the Xist/XIST locus, or its trans regulators, might impact the levels of Xist/XIST expression between individuals, as well as between cells of the same individuals when random XCI is taken into account. Given our findings, Xist/XIST RNA levels are therefore likely to be critical for the establishment and maintenance of sex-specific regulatory programs that when misregulated may lead to phenotypic consequences including sex-biased diseases.

Although previous work in human cells has indicated that XIST might be able to induce some degree of gene silencing in adult cells, these studies did not look into the impact of dialling up Xist levels to induce silencing of facultative and even constitutive escapees. In one study, the ectopic induction of a human XIST cDNA from an autosome in a cancer cell line led to repression of a flanking reporter gene^69^ and in another study, XIST was shown to re-silence genes that had been partially re-activated on the X chromosome in adult human B cells, if XIST was first switched off and then back on using reversible CRISPR interference^49^. Here we directly show that escapees which have either lost a repressive chromatin state (facultative escapees) or never acquired it (constitutive escapees) during differentiation, are still fully susceptible both to Xist-SPEN mediated transcriptional repression in differentiated cells and, in the case of facultative escapees, to the subsequent acquisition of stable epigenetic marks, such as DNA methylation, over time. Thus, although the prevailing view has been that Xist RNA can only act in adult cells via its influence on the Polycomb machinery to reinforce maintenance^12^, our work demonstrates that Xist’s influence on escapees in differentiated cells is via direct SPEN-mediated gene silencing.

Our study also provides important new insights into the interplay between the 3D topology of the Xi and the transcriptional activity of escapees. We show that TAD-like domains encompassing clusters of escapees on the Xi, as previously described in Giorgetti et al 2016^57^, are in fact a direct consequence of gene activity and open chromatin as they are eliminated by gene repression and reappear upon re-establishment of transcription. We show that this reversibility in 3D chromatin organisation is not simply a consequence of higher levels of Xist RNA coating the Xi, but strictly depends on Xist’s SPEN-mediated silencing function. Xist RNA is therefore capable of driving the structural reorganisation of the X chromosome only by inducing gene silencing, and this feature is not limited to early development. Whether the presence of TAD-like structures on the Xi facilitates the escape or reactivation of genes during development or later on in adult tissue is not yet clear. However, our findings show that increased levels of Xist RNA in post-XCI cells can overrule this layer of epigenetic regulation.

Our work also reveals distinctive differences between facultative and constitutive escapees. Although expression of all escapees is affected by increased Xist RNA levels, constitutive escapees are less susceptible and maintain the capacity to resist full XCI, both *in vitro* and *in vivo*. Constitutive escapees are thought to be highly dosage-sensitive and presumably must need to be protected from complete Xist-mediated silencing not only during development, when XCI is first initiated, but also in adult tissues, as any increase in Xist RNA levels could potentially lead to their inactivation with deleterious effects on tissue homeostasis. Indeed, the dosage regulation of constitutive escapees that have retained a Y-chromosome homolog during the evolution of sex chromosomes is thought to be absolutely critical as these genes are involved in many fundamental functions such as chromatin regulation, protein translation and ubiquitination^70,71^.

Notably, the expression levels of constitutive escapees do get affected by increased Xist RNA and this is likely to occur in adult tissues too, with potential consequences on tissue homeostasis. However, the capacity to ultimately override prolonged Xist upregulation, suggest that these genes are likely to have evolved gene-specific strategies that allow them to avoid the repressive epigenetic machinery brought to the Xi by Xist and SPEN. On the other hand, facultative escape is highly variable and likely not driven by the same evolutionary pressures that ensure biallelic expression of constitutive escapees. In particular, whether facultative escape reflects inefficient gene silencing and/or XCI maintenance, or is actually needed (purposeful) is still not fully understood. In fact, there may be genomic and epigenetic features within clusters of escapees that make these regions of the X chromosome more prone to unstable XCI and reactivation. On the other hand, the fact that it is mostly the same subset of facultative escapees that variably resist XCI or become reactivated on the Xi in different tissues or individuals, suggests that higher expression levels of this subset of genes that cluster together on the X chromosome may confer an advantage to XX individuals in certain physiological contexts or under stressful environmental conditions.

Finally, as prolonged high levels of Xist RNA lead to irreversible inactivation of these genes, we speculate that the capacity of facultative genes to become reactivated and to escape XCI in some contexts may be determined by the levels of Xist RNA to which they were exposed at the onset of XCI. In fact, during embryonic development and ESC differentiation, XCI is not perfectly synchronous^28^ and different cells will be exposed to different levels of Xist RNA^41^, potentially setting a threshold for facultative escapees to remain susceptible to varying Xist RNA levels in adult somatic cells. Accordingly, the modulation of their expression levels following changes in Xist RNA expression may underlie the plasticity of this subset of X-linked genes to allow for fine tuning gene dosage regulation in different conditions. Although further studies are necessary to reveal the exact consequences of varying Xist RNA levels to tissue homeostasis, having unveiled the role of Xist RNA in tuning the dosage of escapees provides the basis to ultimately explore the role of this non-coding RNA in disease contexts, and to define the impact of variation in the expression of specific escapee genes on pathological phenotypes, both as biomarkers but also with the perspective of therapeutic strategies that control specific X-linked gene dosage.

## METHODS

### Cell Culture

Mouse ES cells TX1072 (*Mus musculus castaneus* (Cast/Eij) x *Mus musculus domesticus* (C57BL/6)) have been previously derived in the lab ^59^. Cells were cultured on gelatine-coated (0.1 % gelatine in 1X-PBS) cell culture dish in 2i-containing ESC media (DMEM, 15% FBS, 0.1 mM β-mercaptoethanol, 1,000 U/mL−1 leukaemia inhibitory factor (LIF), CHIR99021 (3 µM), and PD0325901 (1 µM)). NPC differentiation and subcloning was performed as previously described ^56^. NPC clones CL30 and CL31 carrying endogenous SPEN alleles tagged with AID-GFP have been previously generated in the lab ^2^. NPCs were cultured on gelatine-coated (0.1 % gelatine in 1X-PBS) cell culture dish in NPC media (N2B27 media supplemented with with FGF2 (10 ng/mL) and EGF (10 ng/mL, both Peprotech)).

### Cell treatments

Xist expression in TX1072 was induced by addition of doxycycline (1 µg/mL) to the NPC media. Culture media supplemented with doxycycline was renewed every 24h. For doxycycline washout, doxycycline-containing media was removed, and cells were refreshed with doxycycline-free culture media. Auxin-mediated depletion of SPEN was achieved by supplementing the culture media with Auxin (Sigma) at the concentration of 500 µM, as previously described ^2^. Auxin-containing media was renewed every 24h.

### RNA fluorescence *in situ* hybridization (FISH)

RNA-FISH on NPCs and preimplantation embryos was performed as previously described ^72,73^. NPCs were dissociated using Accutase (Invitrogen), and attached to Poly-L-Lysine (Sigma) coated coverslips for 10 min. Cells were fixed with 3% paraformaldehyde in PBS for 10 min at room temperature and permeabilized with ice-cold permeabilization buffer (1X-PBS, 0.5% Triton X-100, 2 mM vanadyl-ribonucleoside complex (New England Biolabs)) for 4 min on ice. Coverslips were stored in 70% ethanol at -20 **°**C. Cells were dehydrated in increasing ethanol concentrations (80%, 95%, 100%) and air dried quickly. Probes were prepared from plasmid p510. Probes were fluorescently labelled by nick translation (Abbott). We used dUTP labelled with ATTO-488 green (Jena Bioscience) or Cy5 (Merck). Labelled probes were co-precipitated with mouse Cot-1 DNA (Thermo Fisher) in the presence of ethanol and salt, resuspended in formamide, denatured at 75 **°**C for 8 min and competed at 37 **°**C for 40 min. Probes were hybridised in FISH hybridization buffer (50% formamide, 20% dextran sulfate, 2X SSC, 1 μg/μL BSA (New England Biolabs), 10 mM vanadyl-ribonucleoside complex) at 37 **°**C overnight. The next day coverslips were washed three times for 5 min with 50% formamide in 2X SSC at 42 **°**C, and three times for 5 min with 2X SSC at room temperature. DAPI (0.2 mg/mL was added to the second wash and coverslips were mounted with Vectashield (Vectorlabs). For E3.5 and E4.5 embryos we followed a similar protocol with these modifications: coverslips were incubated in Denhardt’s solution (3X SSC, 0.2 mg/ml BSA, 0.2 mg/ml Ficoll-400, and 0.2 mg/ml polyvinylpyrrolidone (PVP40) in water) for 3 hours at 65 **°**C, followed by incubation in 3:1 methanol/glacial acetic acid solution for 20 min at room temperature and in 0.25% (v/v) glacial acetic acid 0.1 M triethanolamine solution for 10 min at room temperature. The zona pellucida was removed by treatment with acidified Tyrode’s solution. E3.5 embryos were permeabilized for 13 min and E4.5 for 20 min, respectively. Images were acquired with an OLYMPUS iXplore Spin-SR Spinning Disk microscope with 60x or 100x objective. Images were analysed with ImageJ software (Fiji).

### NPC RNA extraction and RNA sequencing

RNA was extracted from > 1M NPCs using the RNeasy kit and on-column DNAse digestion (Qiagen). RNA integrity was measured using the Bioanalyzer Nano Kit. Only high-quality RNA was used for subsequent library preparation using the NEBNext Poly(A) mRNA Magnetic Isolation Module (E7490L) and NEBNext Ultra II Directional RNA Library Prep Kit for Illumina (E7760L, New England Biolabs (NEB), Ipswich, MA, USA) implemented on the liquid handling robot Beckman i7. Obtained libraries that passed the QC step were pooled in equimolar amounts; and the pools were loaded on the Illumina sequencer NexSeq500 or NextSeq2000 and sequenced bi-directionally, generating ∼500 million paired-end reads, each 75 bases long. For the 129/Sv x Cast-EiJ NPCs, RNA was quantified with Qubit RNA Broad Range assay (Invitrogen, Q10211). RNA was sent to Novogene for RNA integrity and purity quality check followed by Eukaryotic strand-specific mRNA (with PolyA-enrichment) library preparation. Libraries were then pooled and sequenced on an Illumina Novaseq 6000 platform for 2x150 bp paired-end reads for 80 million reads total per sample.

### Mouse crosses, embryos collection and whole-embryo RNA sequencing

All experimental design and procedure were performed in agreement with the rules and regulations of the Institutional Animal Care and Use Committee (IACUC) under protocol number 019-03-21EH. Embryos were derived from natural mating between C57BL/6 TX males X^ptet^/Y; R26^rtTA/WT^ (whole embryo RNA-seq) or X^ptet^/Y; R26^rtTA/rtTA^ (RNA FISH) with wild type JF1 females. Embryos were harvested at E2.5 and E3.5 and cultured *in vitro* in G-1 PLUS media (Vitrolife) in the presence of 10 µg/mL Doxycycline for 24 hours. For RNA FISH analysis, approximately half of the collected embryos at each time point were grown in culture medium without Doxycycline. The sex of the embryos was determined either by Xist RNA-FISH or by PCR after RNA-seq libraries preparation. Single embryos were picked and washed three times with transfer buffer (1X-PBS, 0.4% BSA) and transferred into 0.2 ml PCR tubes containing 2μl of lysis buffer (0.7% Triton X-100, 2U/μl RNasin (Promega), 1μl of oligo-dT_30_VN primer (10 μM 5’-aagcagtggttatcaacgcagagtact30vn-3’) and 1 μl of 10 mM dNTP mix (Thermo Fisher). Illumina libraries were prepared by using a modified smart-seq2 protocol [Picelli 2014] using SuperScript IV RT and tagmentation procedure as previously described [Henning 2018]. The RT reaction mix was as follows: 2 μl SSRT IV 5x buffer; 0.5 μl 100 mM DTT; 2 μl 5 M betaine; 0.1 μl 1 M MgCl2; 0.25 μl 40 U/μl RNAse inhibitor; 0.25 μl SSRT IV; 0.1 μl 100 uM TSO, 1.15 μl RNase-free H2O. RT thermal conditions: 52 **°**C 15 min, 80 **°**C 10 min. cDNA was generated using 16 PCR cycles. The cDNA cleanup (0.6x SPRI ratio) was carried out omitting the ethanol wash steps and the elution volume was 13 μl of H2O. For tagmentation, the sample input was normalized to 0.2 ng/μl. Obtained libraries that passed the QC step were pooled in equimolar amounts. After library preparation the sex and genotype of each embryo was assessed by PCR for rtTA (rtTA_F, acgccttagccattgagatg, rtTA_R, tctttagcgacttgatgctc); Xist (Xist_F, ggttctctctccagaagctaggaa, Xist_R, tggtagatggcattgtgtattatatg), and Eif2s3y (Eif2s3y_F, aattgccaggttattttcattttc, Eif2s3y_R, agttcagtggtgcacagcaa). Libraries were sequenced at 50 bp paired-end on NextSeq 2000 platform.

### Capture Hi-C

Capture Hi-C was performed as previously described ^62^. Two arrays of biotinylated RNA probes were designed to tile 3 Mb targets on the mouse X chromosome (*Hcfc1-Mecp2* cluster; ChrX:72,590,000 - 75,430,000; *Kdm5c* cluster; ChrX: 150,210,000 - 153,045,000).

### Enzymatic Methyl-seq (EM-Seq)

Genomic DNA (gDNA) from >1M NPC was extracted using column-based DNeasy Blood & Tissue Kit (Qiagen). DNA integrity was tested on a 0.8% agarose gel and high-quality gDNA was used to prepare libraries according to the NEBNext® Enzymatic Methyl-seq Kit following the section for large insert libraries with minor modifications. A total of 55-100 ng of gDNA was used per library, including a spike-in of pUC and Lambda DNA as a control for methylation efficiency. Samples were fragmented using the Covaris S2 System to achieve an average fragment size of 350-400 bp and barcoded using 5 PCR cycles for 100 ng input and 6 PCR cycles for 55 ng input. The obtained libraries were pooled in equimolar amounts and sequenced at 100 bp paired-end on a NextSeq 2000 platform.

### Targeted capture methylation sequencing (EM-Cap)

Genomic DNA (gDNA) samples used for EM-Seq were used for EM-Cap as previously described ^65^ with minor adjustments. For the final library amplification the cycle number was increased to 14. The same arrays of biotinylated RNA probes designed for the Capture Hi-C experiments were used.

### ATAC-seq

ATAC-seq was performed following the Omni-ATAC protocol ^74^, that has been shown to reduce the number of mitochondrial reads while increasing the signal-to-noise ratio. 25,000 cells were collected, lysed for 3 min on ice in lysis buffer (10mM Tris-HCl pH.5, 5M NaCl, 1M MgCl_2_, 0.1% NP-40, 0.1% Tween-20, 0.01% Digitonin), washed in wash buffer (10mM Tris-HCl pH.5, 5M NaCl, 1M MgCl_2_, 0.1% Tween-20), and spun down at 500 RCF at 4°C for 10 min. Pellets were resuspended in 50 µL of transposition reaction (2X TD buffer, 1X PBS, 0.1% Tween-20, 0.1% Digitonin, 5µL of Illumina Tn5 transposase) and incubated for 30 min at 37°C with 1,000 rpm agitation. DNA was isolated with Zymo DCC5 kit and eluted in 21 µL of elution buffer. DNA samples were initially amplified by 5 cycles of PCR, followed by a variable number of additional amplification cycles estimated by qPCR for each sample. PCR products were purified using the Zymo DCC5 kit and eluted in 20 µL of water. A two-size selection of fragments was performed using 0.5x and 1.3x volume of AMPure XP beads (Beckman Coulter). Libraries were quantified and analyzed using Qubit and Tapestation assays, before preparing equimolar dilutions. Paired-end sequencing was performed on NextSeq 500 (Illumina).

## BIOINFORMATICS

### Allele-specific preprocessing of RNA-Sequencing data

All steps for the preprocessing of RNA-Sequencing data can be reproduced using a nextflow pipeline available at https://github.com/yuviaapr/allele-specific_RNA-seq. Reads were trimmed using trim_galore (v0.6.6). To construct reference genomes for allele-specific mapping, genomes in which known heterozygous variants (https://ftp.ebi.ac.uk/pub/databases/mousegenomes/REL-2112-v8-SNPs_Indels/mgp_REL2021_snps.vcf.gz) were masked by the ambiguous base N were constructed using SNPsplit (*SNPsplit_genome_preparation* script, v0.5.0) and converted to STAR references (STAR v2.5.3a). For the E6, CL30/CL30.7 and CL31/CL31.16, the mm10 genome (GRCm38) was used. For the embryo and 129/Sv x CAST/EiJ cell lines, the mm11 genome was used (GRCm39). Reads were aligned to the N-masked genomes using the options *--sjdbOverhang 99 -- outFilterMultimapNmax 1 --outFilterMismatchNmax 999 --outFilterMismatchNoverLmax 0.06 -- alignIntronMax 500000 --alignMatesGapMax 500000 --alignEndsType*. Reads mapping to the mitochondrial genome were removed and split into parental genotypes using SNPsplit and known heterozygous variants. Gene-level read counts for each haplotype and without allelic resolution were derived using featureCounts (v2.0.1).

### Computation of allelic ratios in RNA-Seq for escapee definition

Allele-specific expression per gene was calculated from genotype-assigned read counts. First, lowly expressed genes with an average allelic read count < 10 were excluded (summing both haplotypes). Escape of an X-linked gene was generally quantified as the fraction of read counts on the inactivated X against the total read count across both active and inactive X (Xi / (Xi + Xa), allelic ratio, also known as d-score). Genes were considered as escaping when the allelic ratio exceeded 0.1 (10% of expression from the inactive X). As the causal gene of XCI, *Xist* was not considered an escapee. A small number of other genes showed allelic ratios > 0.8, which likely represents either strain-specific expression or technical artefacts due to erroneous mapping or SNPs with non-reference genotypes, rather than genuine expression largely restricted to the Xi. These genes were excluded for subsequent analysis. For all analysis of escape in the E6 cell line, we used the set of 133 genes (134 including *Xist*) which showed an allelic ratio > 0.1 in all untreated replicates. The use of N-masking of the genome should reduce mapping bias at SNP positions. Furthermore, we focus on relative changes (reductions) in allelic ratios, which should be unaffected by technical artefacts.

### Escapee meta-analysis

To obtain a consensus of escaping genes from the literature, we performed a meta-analysis of papers that used genome wide expression profiling to derive escapees. For each study (Supplementary Table 1), we considered the genes defined as escapees or silenced by the experimental and analysis methodology used in that paper. In each study, a gene is therefore considered “escaping”, “silenced” or “not detected”. We then classified genes as “constitutive escapees”, if they were detected in at least 3 and escaped in more than 50% of the studies. We defined genes as “facultative escapees”, if they were escaping in less than 50% of studies or were only detected in less than 3 (but escaped in at least one study). Finally, we defined genes as “silenced”, if they were not escaping in any assayed study.

### Differential allelic imbalance analysis

To test for differential allelic imbalance (differences in escape between conditions), we employed binomial generalized linear models (R package *stats, v4.2.0*). We modelled the number of reads mapping to the inactive X as binomially distributed with the formula (k, n) ∼ 1 + treatment and tested for the significance of a treatment coefficient using a Wald test. P-values were adjusted using the Holm-Bonferroni method.

### Modelling of escape trajectories

We used non-linear parametric regression to fit an exponential decay curve to the allelic ratios after *Xist* overexpression. Specifically, we used the *nls* function (R, *stats*) to fit the allelic ratios *r = Xi / (Xi + Xa)* with the exponential decay function *r ∼ a * exp(- k * t) + b*, where *t* represents the number of days of doxycycline treatment, *a* is a scaling parameter of the decay, *b* allows for an asymptotic offset and *k* specifies the decay constant. In particular, *k* relates to the speed (half-life) of silencing loss by *t1/2 = - ln(0.5)/k* and *b* quantifies whether there is residual escape after prolonged *Xist* overexpression. We also fit the same model without *b*, to test which genes showed evidence for residual escape as opposed to full silencing. After excluding all genes with a regression R∧2 < 0.3 for both fits, we used the Bayesian Information Criterion to classify whether genes showed residual escape.

### Definition of escapee clusters

To define locally close groups of escapees on the X-chromosome, we iterated through all escapees on the X, grouping genes into one “cluster” if the gene start position was within 100kb of each other, and splitting whenever this was not the case. This approach partitioned the escapee set into 11 groups of 3 or more genes (with 3 - 14 escapees) and 57 singles or pairs.

### Differential expression analysis

We used DESeq2 (R, v1.36.0) to test both haplotype-specific and total expression counts for differential expression using standard settings, using the formula ∼ Treatment and adjusting p-values using the Benjamini-Hochberg correction.

### Allele-specific analysis of DNA methylation data

Processing of DNA methylation was performed using the methylseq nextflow pipeline with the --emseq option (*v2.3.0*, NextFlow *v22.10.6*). Reads were trimmed using TrimGalore! (*v0.6.7*) and aligned to the same N-masked genome (GRCm38) using bismarck (*v0.24.0*), and used SNP_split (v0.5.0) to assign mapped reads to parental genomes, similar to the RNA-Seq data. We finally used bismark_methylation_extractor from the bismarck software to generate base-level methylation counts for both haplotypes. We retrieved genomic coordinates of CpG islands (http://genome.ucsc.edu/cgi-bin/hgTrackUi?g=cpgIslandExt), gene bodies and promoters (2kb up-to 200bp downstream of transcription start sites) (*EnsDb.Mmusculus.v79*, *v2.99.0*). To quantify DNA methylation in promoters, we used regions of accessible chromatin defined by ATAC-Seq data in matched cell lines (see below) if they overlapped with annotated promoter regions. We quantified average methylation values across per-base methylation values b = methylated / (methylated + unmethylated) in the regions of interest.

### Allele-specific Capture Hi-C analysis

Allele-specific Capture Hi-C analysis was performed using the Hi-C Pro pipeline (v2.11.4) ^75^ as previously described ^62^. Paired-end reads were first independently aligned using bowtie2 to the same N-masked genome reference. Read pairs were then assigned to either C57BL-6J or CAST-EiJ genomes and allele-specific BAM files for downstream analysis are generated. Low-quality reads, multiple hits, singletons and read pairs which do not map to the same genotype were then removed. Further filtering for valid interactions, which excludes reads outside of the captured region, was performed and only valid pairs were used to build the normalized contact maps at a 10 kb resolution with the cooler package (v0.8.9). Insulation scores were calculated using the cooltools package (v0.3.2) ^76^ and boundaries were deduced from it a the local minimas. All data was visualized using pyGenomeTracks ^77^.

### Allele-specific ATAC-seq analysis

All steps for the preprocessing of ATAC-seq data can be reproduced using a nextflow pipeline available at https://github.com/yuviaapr/allele-specific_ATAC-seq. Briefly, initial quality checks of the raw FASTQ files were performed using FastQC (v0.11.9). Reads were trimmed using Trim Galore (v0.6.3). Reference genome for allele-specific mapping, was constructed as described for the allele-specific RNA-seq analysis, and used to generate a bowtie index. Paired-end reads were aligned using bowtie2 (v2.3.4.1) to the N-masked genome index with the following parameters: *--very-sensitive -X 2000*. Duplicates were removed using Picard Tools (v2.20.8) before downstream analyses. Peaks were called using MACS2 (v2.2.7.1) using the following parameters: *-g mm --buffer-size 100 -q 0.01 --keep-dup all --min-length 100 --format BAMPE --nomodel*. Consensus peaks were generated using the “*dba.peakset*” function from the DiffBind Bioconductor R package (v3.8.4), with the requirement that peaks should be present in at least two BAM files. Blacklist regions (from ENCODE for *BSgenome.Mmusculus.UCSC.mm10*) were removed using the *dba.blacklist* function, and the raw counts for the resulting consensus peak set were generated using the *dba.count* function using *summits=FALSE*. Peak counts were normalised using the *dba.normalize* function, with default settings. Differential accessibility analysis between the split BAM files coming from the Xi and Xa was performed using the dba.analyze function with *methods=DBA_DESEQ2* (calling the DESeq2 Bioconductor R package (v1.38.3) in the background). Peaks were annotated to their nearest gene using the ChIPseeker Bioconductor R package (v1.34.1) using the mm10 genome from the TxDb.Mmusculus.UCSC.mm10.knownGene Bioconductor R package (v3.10.0).

### Processing of embryo RNA-Seq data

The embryo RNA-Sequencing data was pre-processed as described above (**Allele-specific preprocessing of RNA-Sequencing data**), using the SNPs between C57BL/6 and JF1 mice (https://ftp.ebi.ac.uk/pub/databases/mousegenomes/REL-2112-v8-SNPs_Indels/mgp_REL2021_snps.vcf.gz). To confirm the annotated genotypes of the embryos, the rtTA transgene was included in the reference genome, and any embryo with >50 mapped transgene reads was considered rtTA-positive. For developmental staging, principal component analysis (*prcomp*, stats *v4.2.0*) was performed on total read counts after size-factor normalization (*estimateSizeFactors*, DESeq2, *v1.36.0*) and subsetting on highly variable genes (getTopHVGs, scran, *v1.24.1*). The first principal component separated embryo stages as determined by morphological staging and was used as a developmental pseudotime. For the allelic analysis, genes were considered escapees if they showed an allelic ratio > 0.1 in at least 50% of embryos. Genes were annotated into escape categories as in the meta-analysis and classified as “embryo-specific” if they did not escape in E6 and/or were not annotated as escapees in the meta-analysis.

## Data availability

RNA-seq, Capture Hi-C, ATAC-seq and Methyl-seq data generated in this study have been deposited in the Gene Expression Omnibus under accession number GSEXXXXXX

## Code availability

Code available upon request

## ACKNOWLEDGMENTS

This work was funded by an ERC Advanced Investigator award to E.H. (XPRESS - AdG671027), the European ITN Innovative and Interdisciplinary Network “ChromDesign”, under the Marie Skłodowska-Curie Grant agreement 813327 to E.H., a Marie Skłodowska-Curie Actions Individual Fellowship to A.L. (IF-838408); Core funding from the German Cancer Research Center (O.S & D.T.O) and the European Molecular Biology Laboratory (O.S), the European Research Council (grant agreement IDs 810296 / DECODE to O.S. and 788937/ CTCFStableGenome and 615584 / EvoGeneticsTFBinding to D.T.O), Wellcome Investigator Award (202878_Z_16_Z to D.T.O.), as well as the Bundesministerium für Bildung und Forschung Germany, project MERGE, Förderkennzeichen 031L0174C (to O.S.). H.Y.C. is an Investigator of the Howard Hughes Medical Institute and he is supported by NIH RM1-HG007735. We thank T. Pollex for critical and thoughtful reading of the manuscript, P. Ginno for discussion and members of the Heard, Stegle and Odom lab for discussion. We also thank Vladimir Benes and the Genomics Core Facility at EMBL (Heidelberg), for support and assistance.

## AUTHOR CONTRIBUTION

A.L. and E.H. conceived the study. A.H. performed the experiments trained by A.L. and with technical support from I.R. unless stated otherwise. J.P. processed and analysed all presented data unless stated otherwise. C.P. and A.L. generated the NPC line and performed RNA-FISH and some of the RNA-seq experiments. L.V. prepared RNA-seq and Capture Hi-C libraries, F.J. prepared EM-seq and EM-cap libraries. Y.A.P. and L.C. performed the ATAC-seq experiments and compiled the reference list of escapees. Y.A.P. processed the ATAC-seq data and N. Servaas performed the analysis. N. Servant analysed the Capture Hi-C data. A.L. and E.K. performed the in vivo experiments. C.L. prepared RNA-libraries for the 129/Sv x Cast EiJ NPCs. O.S. and D.O. supervised J.P. and provided feedback throughout the project. A.L. and E.H. supervised the work and wrote the manuscript with input from J.P., D.O., O.S. and all authors.

