## Supplementary Figures for "Escape from X inactivation is directly modulated by levels of Xist non-coding RNA" (from another version of this preprint: version 1, 2024-02-24)

**Extended Data Fig.1**

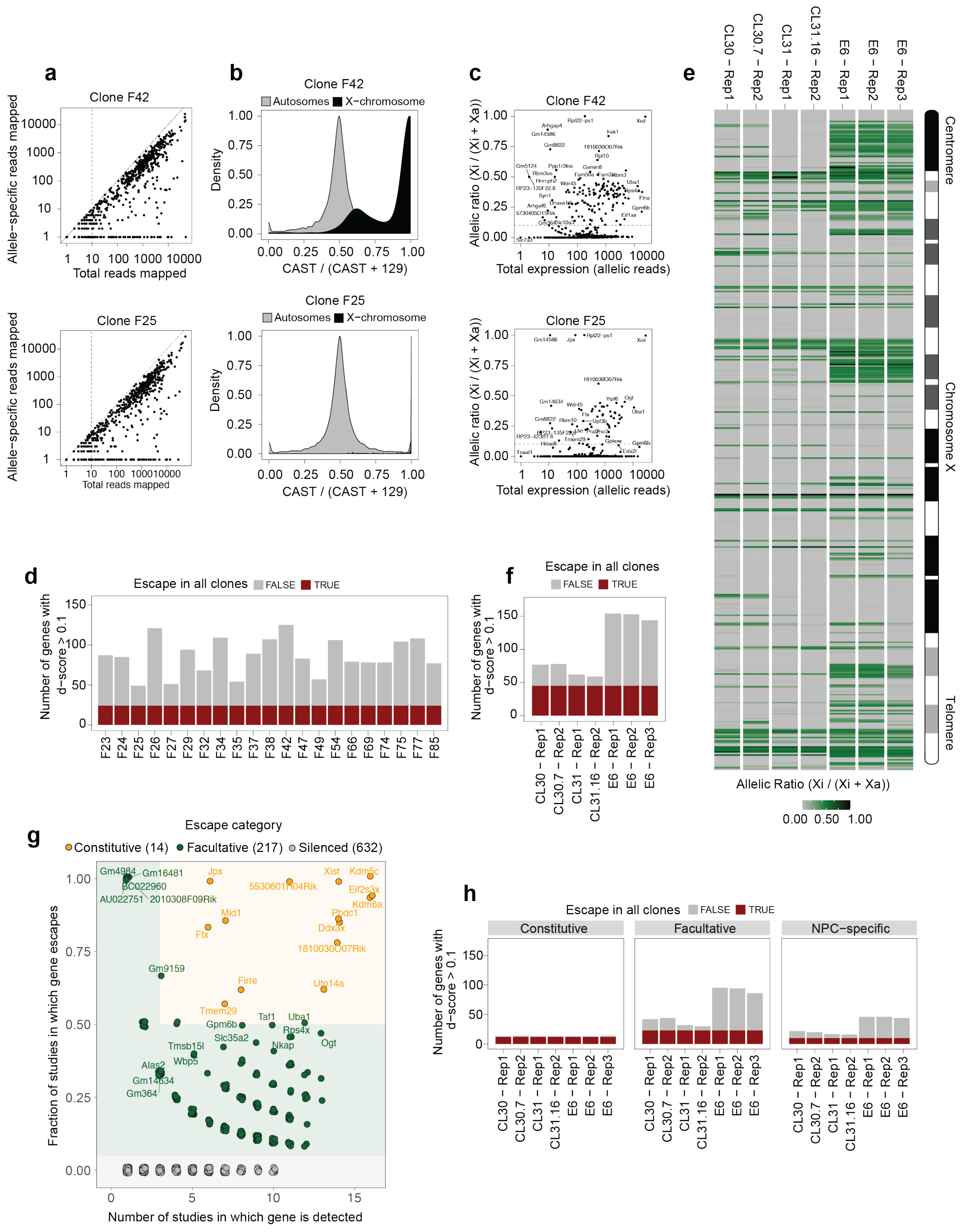

**Extended Data Fig.1: Quantification of escape across neural progenitor cell lines**

**a,** Scatterplots showing the total number of reads mapping to each gene (x-axis) and the number of allele-specific RNA-Seq reads (y-axis) in two out of 21 clonal F1 hybrid neural progenitor cell lines (129/Sv x Cast/Eij) cell lines (F42 and F25). The diagonal represents equal values. **b,** Density plot showing the distribution of allelic ratios (CAST / (CAST + 129); Xa / (Xi + Xa)) for the X-chromosome (black) and all autosomes (grey) F42 and F25. Those cell lines were chosen to represent high (F42) and low (F25) general escape. **c,** Scatterplot showing the total number of reads per gene against their allelic ratio (Xi / (Xi + Xa)). Most genes show an allelic ratio of close to 0 (no expression from the Xi), while a subset shows higher allelic ratios (escape from XCI). **d,** Barplot quantifying the number of escapees (allelic ratio > 0.1) across the 129 x CAST clones. The red bar indicates the number of genes escaping in all 21 clones (24). **e,** Schematic of the mouse X chromosome and heatmap showing the allelic ratios of expressed X-linked genes in 3 NPC clones. CL30.7 and CL31.6 are sub-clones of CL30 and CL31 (Dossin et al. 2020). Three replicates are shown for clone E6. **f,** Quantification of the number of escapees in F1 hybrid NPC clones (C57BL/6 x Cast/Eij) by allele-specific RNA-seq as in **d**. **g,** Classification of X-linked genes in three categories based on their XCI status as described in 21 studies spanning multiple cell type and developmental contexts: (i) constitutive escapee (escape XCI in >50% of the studies); (ii) facultative escapees (variably escaping XCI in different studies and in at least one of the considered studies); and (iii) silent genes (escape in no study). **h,** Classification of the escapees identified in this study in clones CL30, CL31 and E6 according to the categories defined in **g**. As in **d**, the red bar indicates escape in all cell lines and replicates.

### Extended Data Fig. 2

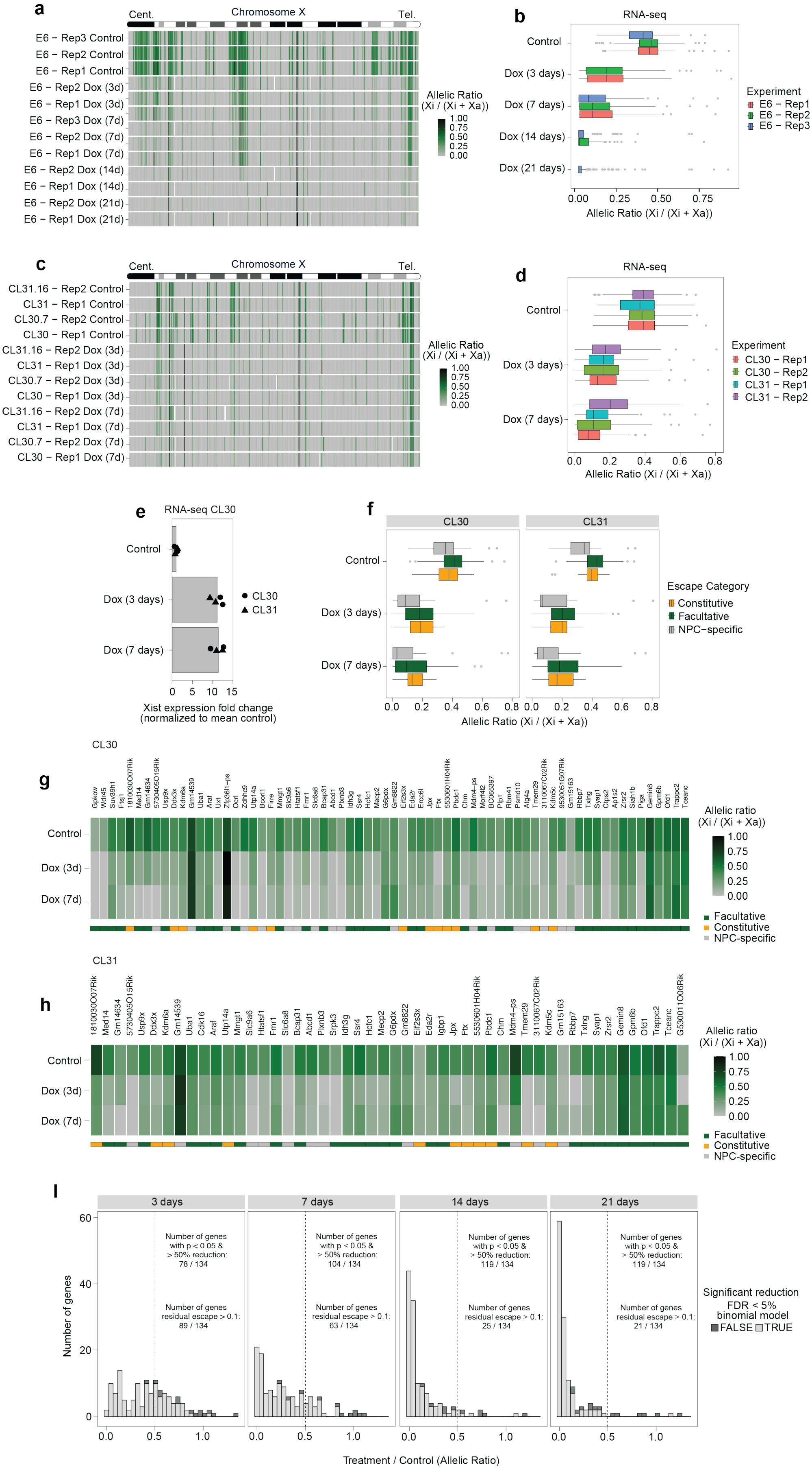

**Extended Data Fig. 2: Reduction of escape during *Xist* treatment across clones and replicates**

**a,** Schematic of the mouse X chromosome and heatmap showing X-linked transcript allelic ratios for expressed genes in untreated clone E6 and after 3, 7, 14 and 21 days of doxycycline treatment. Data relative to 1 to 3 replicates per time point is shown. **b,** Box plot showing the changes in allelic ratios of escapee distributions across the time course of doxycycline treatment for E6 clones. **c,** Same as **a,** for clones CL30 and CL31 as well as subcloned lines. **d,** Same as **a,** for clones CL30 and CL31 as well as subcloned lines. **e,** RNA-seq data showing the fold change of *Xist* expression compared to untreated cells across the time course of doxycycline treatment. **f,** Box plot showing the changes in allelic ratios for different categories of escapees across the time course of doxycycline treatment for CL30 and CL31. Shown is the average for the primary and subcloned cell lines. **g,** Heatmap showing the allelic ratios of escapees identified in clone CL30 as shown in **c**. Escapees are assigned to three different categories as described in **Fig. S1.** The escape category for each gene is indicated below the heatmap. **h**, same as **g,** for clone CL31. **i,** Barplots quantifying the decrease in escape upon doxycycline treatment. Shown is the allelic ratio (AR) after treatment normalised on the untreated sample for each gene (ARXist / ARuntreated). Additionally, genes are stratified based on whether a significant decrease in allelic ratio was observed (binomial linear model, **Methods**) and whether they retained an allelic ratio > 0.1.

### Extended Data Fig. 3

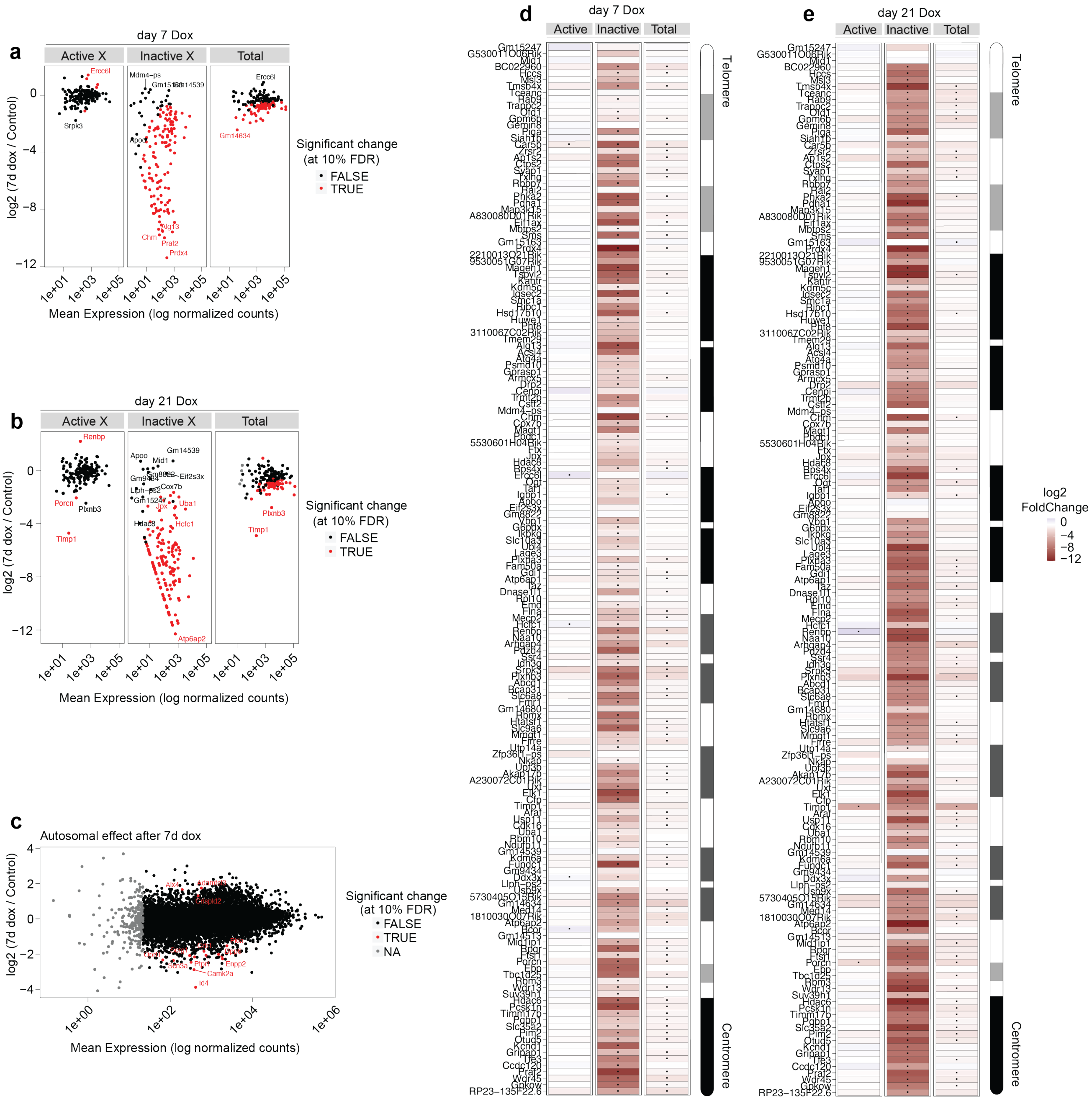

**Extended Data Fig. 3: Allele-specific and autosomal differential expression analysis**

**a,** MA-plots showing the results of a differential expression analysis after 7 days of doxycycline treatment for X-linked genes considering active counts, inactive counts and all reads (including those not assignable to an allele). p-values are computed using DESeq2 and FDR-adjusted. **b,** As **a**, after 21d of treatment. **c,** MA-plot showing differential expression analysis of autosomal genes after 7 days of doxycycline treatment. **d,** Heatmap showing the fold changes of X-linked genes after 7 days of doxycycline treatment as summarised in **a** for each gene individually. **e,** As **d**, after 21 days of doxycycline treatment.

#### Extended Data Fig. 4

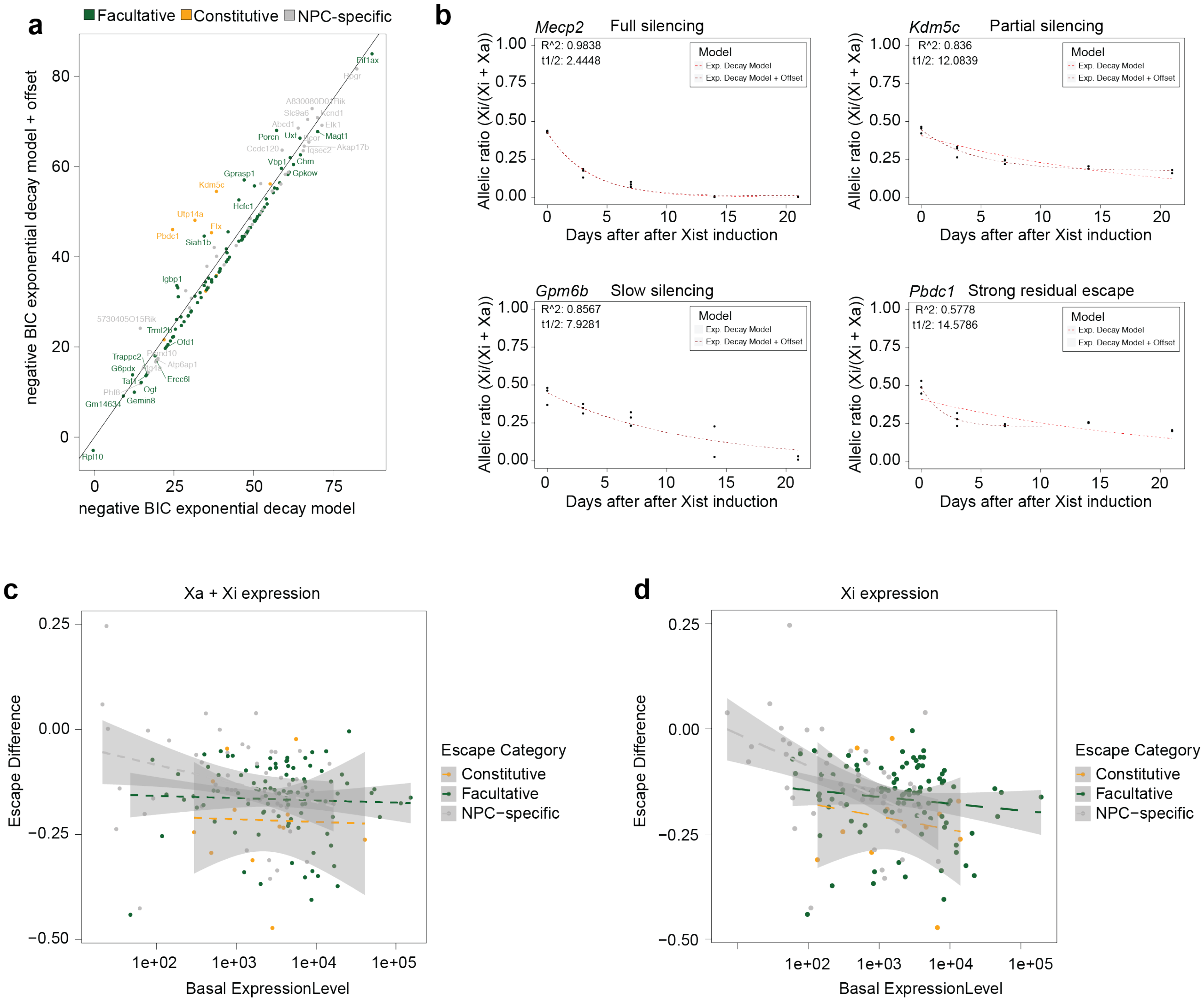


**Extended Data Fig. 4: Supplement to modelling of decay curves**

**a,** Scatterplot showing the negative Bayesian Information Criterion to compare model fits between the exponential decay model and the exponential decay model with offset. The diagonal indicates equal values and for genes above the diagonal the data is better explained by a non-zero offset parameter. **b,** Example model fits for different genes, including the R2 goodness-of-fit statistic for the full silencing model, and the silencing half-life. Genes are shown to represent the spectrum of response curves, including complete and fast silencing (*Mecp2*), complete but slow silencing (*Gpm6b*), partial silencing (*Kdm5c*) and fast silencing but strong residual escape (*Pbdc1*). **c,** Scatterplot showing that the magnitude of change in escape is not a function of gene expression level across Xi and Xa. The escape difference represents the difference in allelic ratio between 7d of treatment and untreated cells. **d,** As (**c**), but comparing expression differences against expression level on the Xi only.

**Extended Data Fig. 5**

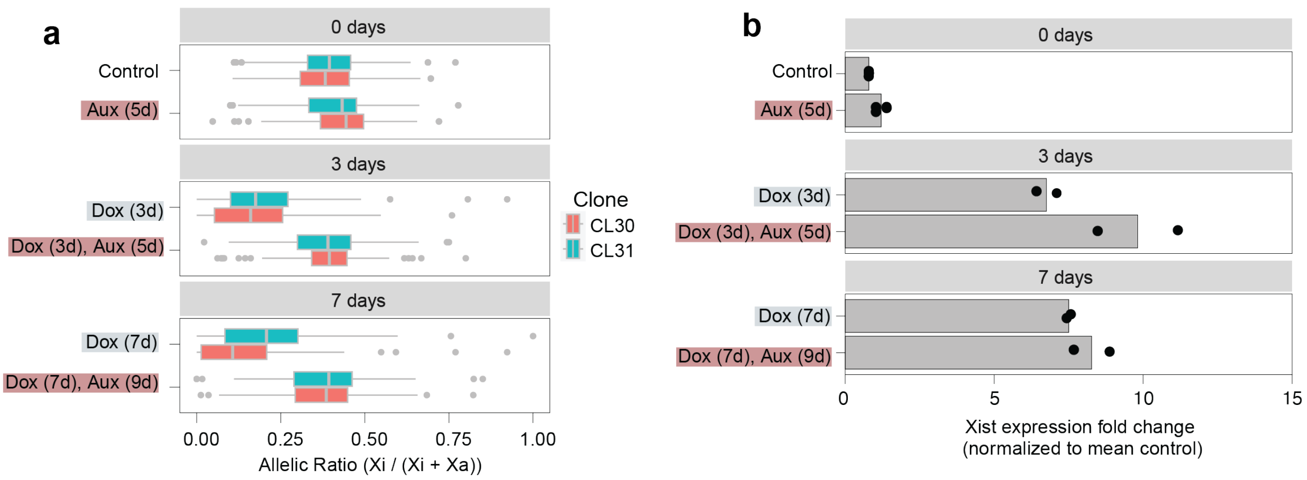


**Extended Data Fig. 5: Supplement to SPEN-depletion experiments**

**a,** Box plot showing the changes in allelic ratios of escapees across the time course of doxycycline and auxin treatment. Data from sub-clonal lines CL30.7 and CL31.16 is shown. **b,** RNA-seq data showing the fold change in Xist expression compared to untreated cells following doxycycline treatment for 3 and 7 days and in combination with auxin treatment for 5 and 9 days.

#### Extended Data Fig. 6

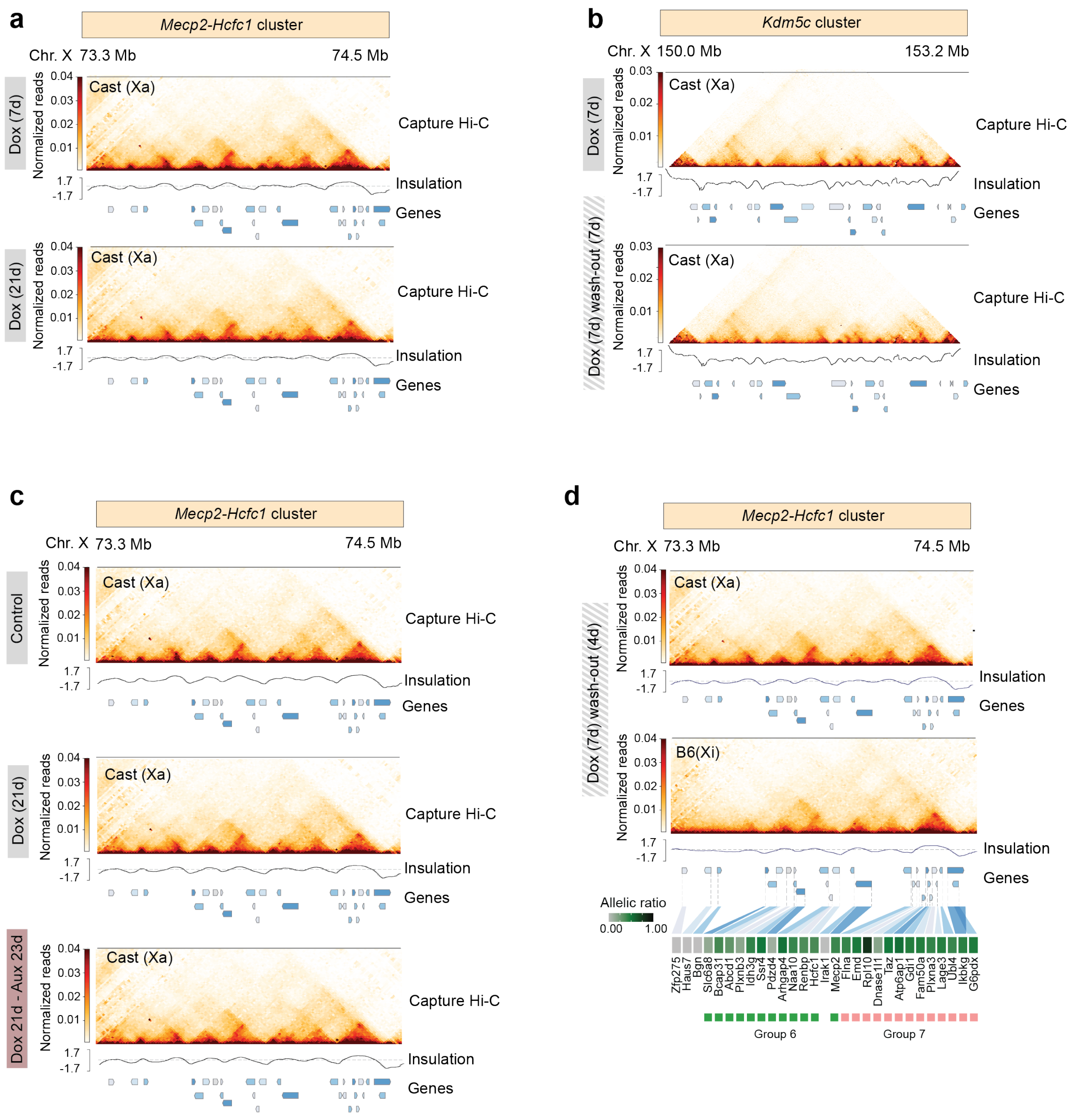


**Extended Data Fig. 6:**

**a-b** Capture Hi-C interactions and insulation score at **(a)** *Mecp2-Hcfc1* and **(b)** *Kdm5c* clusters in clone E6 on the Xa after **(a)** 7 and 21 days of doxycycline treatment and **(b)** 7 days of doxycycline treatment and 4 days of doxycycline wash-out. **(c)** Capture Hi-C interactions and insulation score at the *Mecp2-Hcfc1* cluster in clone CL30.7 on the Xa in control condition, after 21 days of doxycycline treatment, and after 23 days of auxin treatment in combination with 21 days of auxin treatment. All Capture Hi-C data is shown at 10-kb resolution. **d,** Capture Hi-C interactions and insulation score at the *Mecp2-Hcfc1* cluster on the Xa and Xi after 7 days of doxycycline treatment followed by 4 days of doxycycline wash-out. Capture Hi-C data is shown at 10 kb resolution. Heatmaps showing allelic ratios for 29 X-linked genes included in the captured region are shown. Escapees belonging to groups 6 (green) and 7 (pink) within the *Mecp2-Hcfc1* cluster are highlighted.
